# Plasticity-induced actin polymerization positions endocytic compartments at the bases of stimulated spines

**DOI:** 10.64898/2026.10.08.756800

**Authors:** V. C. Wong, D. Walpita, Z. J. Liu, E. K. O’Shea

## Abstract

The endocytic pathway plays key roles in facilitating and regulating synaptic plasticity, but how its components are organized at spines undergoing plasticity is not fully understood. Here we use live-cell imaging and single-particle tracking to examine the localization of endosomes and lysosomes in response to plasticity-inducing stimuli. We find that endosomes and lysosomes are positioned and retained near the bases of stimulated spines after the induction of structural plasticity in a manner dependent on actin polymerization in the dendritic shaft. Spines with endocytic compartments near their bases exhibit greater glutamate receptor turnover and more sustained enlargement after stimulation than spines lacking them. These results suggest that actin polymerization in the dendritic shaft acts as a hub to organize critical cellular machinery at the bases of spines undergoing synaptic plasticity.

## Introduction

Synaptic plasticity – the modification of synaptic strength or efficacy in response to neuronal activity – is regarded as the primary cellular correlate of learning and memory (Citri and Malenka, 2008). Plasticity can be characterized by changes to synaptic structure (i.e., structural plasticity; Nakahata and Yasuda, 2018) and the composition of proteins in synapses that result in the modulation of excitatory postsynaptic currents (i.e., functional plasticity; Magee and Grienberger, 2020). The endocytic pathway – the organelles and machinery responsible for recycling and degrading proteins internalized from the cell membrane – supports biochemical changes inside spines during synaptic plasticity (Wilkinson and Lin, 2004; Kelly et al., 2011; Camblor-Perujo and Kononenko, 2022; Grochowska et al., 2022). Critically, endosomes maintain a reservoir of glutamate receptors that can be inserted into synapses to strengthen connections when plasticity is induced (Park et al., 2004; Park et al., 2006; Kneussel and Hausrat, 2016). How the endocytic pathway is spatially organized to support these functions, however, is not well understood.

α-amino-3-hydroxy-5-methyl-4-isoxazolepropionic acid receptors (AMPARs) are ionotropic glutamate receptors that are rapidly trafficked to and from synapses to modulate excitatory postsynaptic currents (Chater and Goda, 2014). AMPARs on the cell membrane are internalized via clathrin-dependent endocytosis (Man et al., 2000; Carroll et al., 2001) and then sorted inside early endosomes into either recycling or degradative pathways, depending on synaptic activity (Ehlers, 2000). In hippocampal neurons, the activation of N-methyl-D-aspartate receptors (NMDARs) drives AMPAR internalization, after which receptors are recycled and reinserted into the plasma membrane, whereas AMPARs internalized in the absence of NMDAR activation are degraded (Ehlers, 2000).

Several observations suggest that endosomes and lysosomes are targeted to specific locations and not randomly distributed. Endocytic compartments are transported along the length of dendrites on microtubules in anterograde and retrograde directions (Yap and Winckler, 2022) and can be found both inside and at the bases of spines (Esteves da Silva et al., 2015; Goo et al., 2017; Yap et al., 2018). For example, recycling endosomes can be found inside spine heads, where they regulate spine structure and the concentrations of scaffolding proteins and glutamate receptors (Esteves da Silva et al., 2015; Bowen et al., 2017). Lysosomes are also found throughout the dendritic shaft, often associated with F-actin patches (van Bommel et al., 2019), and sometimes inside spine heads (Goo et al., 2017). The distribution of lysosomes is correlated with the internalization of membrane proteins near synapses and, importantly, lysosomes can be found paused at the bases of spines undergoing synaptic activity (Goo et al., 2017).

Despite these advances, it remains unclear whether a general mechanism targets components of the endocytic pathway (early/sorting, late, and recycling endosomes, in addition to lysosomes) to spines undergoing plasticity. We used fluorescence timelapse imaging and single-particle tracking (SPT; Sergé et al., 2008) combined with hidden Markov modeling with Bayesian model selection (HMM-Bayes; Monnier et al., 2015; Wong et al., 2025) to demonstrate that endocytic compartments are stably positioned near the bases of spines undergoing structural plasticity. The stable positioning of endosomes and lysosomes is correlated with, and requires, actin polymerization in the dendritic shaft surrounding the bases of stimulated spines. Finally, using a pulse-chase protocol with cell-impermeable Janelia Fluor dyes (Xie et al., 2017; Jonker et al., 2020), we find that positioned endosomes and lysosomes predict the magnitude of AMPAR turnover at stimulated spines. These results demonstrate that F-actin in the dendritic shaft acts as an organizational hub to recruit components of the endocytic pathway to activated spines.

## Results

### Endosomes and lysosomes are positioned near the bases of spines when structural plasticity is induced

Although previous studies have established roles for the endocytic pathway during synaptic plasticity (Kneussel and Hausrat, 2016), where endosomes and lysosomes (hereafter abbreviated as ELs) are positioned relative to activated spines is not fully characterized. We first asked whether ELs are recruited to spines undergoing synaptic plasticity. To track the location of ELs in live cells, we expressed fluorescent protein (FP)-tagged Rab4, Rab7, Rab11, or LAMP1 in cultured rat hippocampal neurons (**Figure 1A**). These proteins play key roles in regulating the functions of early/sorting, late, and recycling endosomes and lysosomes, respectively (Chen et al., 1985; van der Sluijs et al., 1992; Ullrich et al., 1996; Vitelli et al., 1997; Somsel Rodman and Wandinger-Ness, 2000; Eskelinen, 2006). Tagged versions of these proteins are functional (Bucci et al., 2000; Sönnichsen et al., 2000; Knödler et al., 2010; D’Souza et al., 2014; Goo et al., 2017) and reflect the localization of the native, untagged proteins (**Figure S1**) and thus are used as markers to image their localization (Bastin and Heximer, 2013). When FP-tagged Rabs or LAMP1 are expressed, ELs appear as bright puncta under fluorescence microscopy that can be readily identified and tracked (**Video 1**). We utilized a single-photon (1p) 4-methoxy-7-nitroindolinyl-caged-L-glutamate (MNI) uncaging protocol (**Figure 1B**; Ellis-Davies, 2007) to induce structural long-term potentiation (sLTP; Matsuzaki et al., 2004) at selected dendritic spines and examined the motion of Rab^+^ endosomes or LAMP1^+^ lysosomes. We use sLTP to induce plasticity because we and others have demonstrated that this protocol stimulates the expansion of spine volume and the rapid recruitment of AMPARs to targeted spines, important markers for structural and functional plasticity, respectively (Matsuzaki et al., 2001; Matsuzaki et al., 2004; Huganir and Nicoll, 2013; Bosch et al., 2014; Wong et al., 2024). Critically, this protocol can be used to induce plasticity at a single spine (Matsuzaki et al., 2004; Wong et al., 2024), enabling us to determine whether ELs are recruited to the stimulated spine.

**Figure 1.**
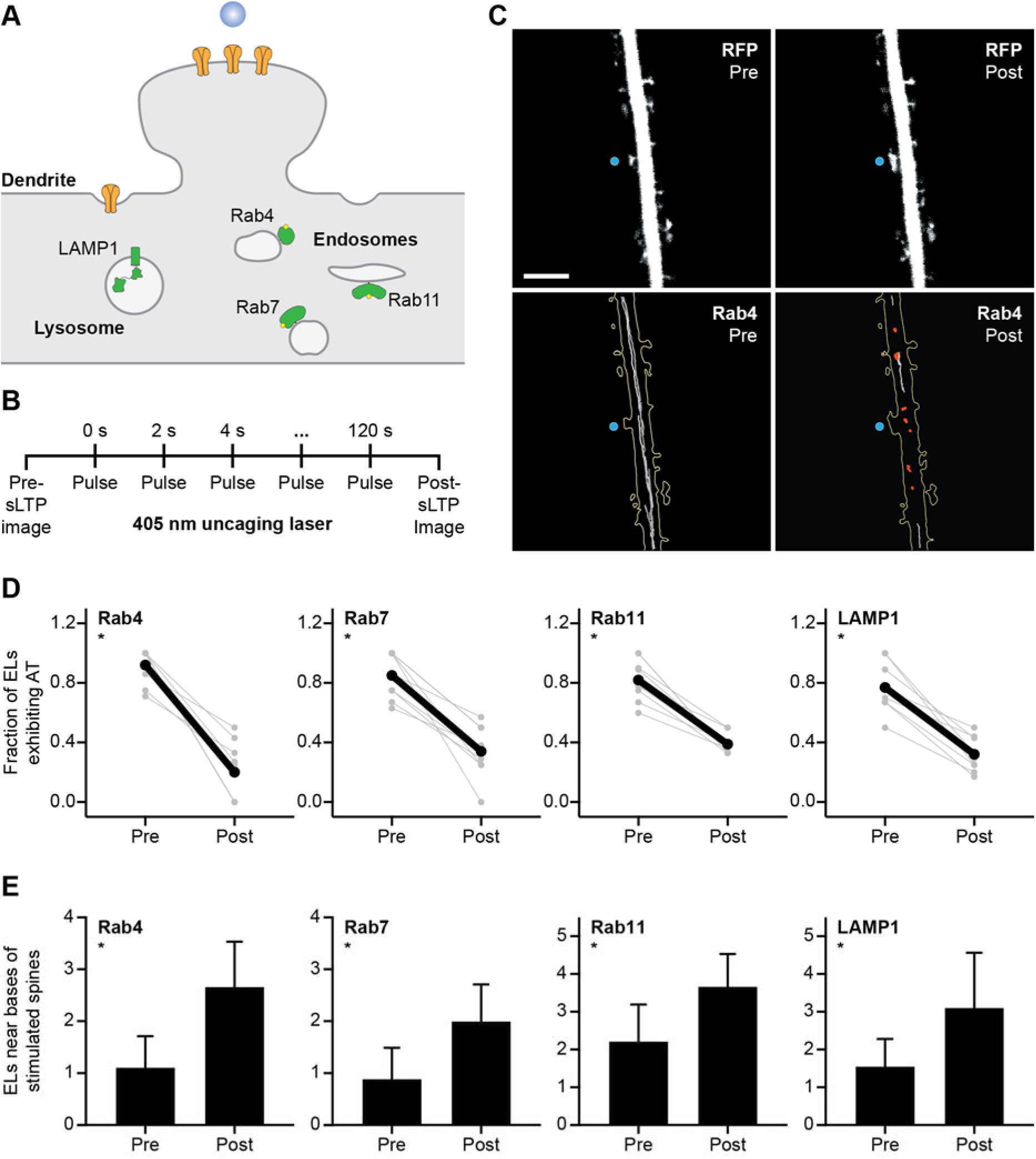
Endosomes and lysosomes are positioned near the bases of stimulated spines when structural plasticity is induced. (A) Diagram depicting the recruitment of endosomes and lysosomes to the base of a spine undergoing plasticity. Fluorescent protein (FP)-tagged Rab4, Rab7, Rab11, and LAMP1 are used as markers to identify ELs. Yellow receptors represent AMPA receptors. Blue dot represents plasticity-inducing stimulus. (B) Single-photon (1p) 4-methoxy-7-nitroindolinyl-caged-L-glutamate (MNI) uncaging protocol used to stimulate structural long-term potentiation (sLTP). A 405 nm laser is fired 1 µm from a spine head for 2 minutes at 0.5 Hz (see **Methods**). (C) RFP (top): representative spinning disk confocal images of a dendritic spine expressing tdTomato pre-sLTP (Pre) and post-sLTP (Post). Blue dot indicates MNI uncaging site. Scale bar, 5 µm. tdTomato was used to identify and segment the dendritic spine and shaft. Rab4 (bottom): Rab4-mEmerald^+^ (Rab4^+^) endosome trajectories in the dendrite depicted above (RFP, top) pre- and post-sLTP. Gray trajectories indicate active transport; red trajectories indicate diffusive motion. (D) Fraction of mEmerald-tagged Rab4^+^, Rab7^+^, or Rab11^+^ endosomes, or LAMP1^+^ lysosomes (ELs) exhibiting active transport (AT) pre- and post-sLTP. Each pair of gray dots represents the fraction of ELs exhibiting AT in a dendrite pre- and post-sLTP for one stimulation experiment. Black dots represent the mean fraction of ELs exhibiting AT pre- and post-sLTP. Rab4, *p=0.0039; Rab7, *p=0.0039; Rab11, *p=0.0039; LAMP1, *p=0.0039. Significance determined by Wilcoxon signed-rank test with Holm–Šídák correction across four EL markers. N = 9 sLTP stimulation experiments for each marker; for each experiment, the motion of ELs in the dendrite was imaged before and after sLTP (see **Methods**). A new neuron was selected for each sLTP stimulation experiment; a maximum of three spines (i.e., neurons) was stimulated per neuronal culture. (E) Number of ELs that appear within a 4 µm segment of dendrite below stimulated spines (see **Figure S2**) pre- and post-sLTP. Bars and error bars represent mean and standard deviation, respectively. Rab4, *p=0.0039; Rab7, *p=0.0156; Rab11, *p=0.0078; LAMP1, *p=0.0234. Significance determined by Wilcoxon signed-rank test with Holm–Šídák correction across four EL markers. N = 9 sLTP stimulation experiments for each marker; for each experiment, the motion of ELs in the dendrite was imaged before and after sLTP, and the number of trajectories within a 4 µm segment of dendrite below the stimulated spine was counted for each condition (see **Methods**). A new neuron was selected for each sLTP stimulation experiment; a maximum of three spines (i.e., neurons) was stimulated per neuronal culture.

We first tested whether inducing plasticity would alter the motion of Rab4^+^ endosomes near stimulated spines (**Figure 1C**; **Video 2**). sLTP triggers a significant expansion in spine area at the targeted spine (**Figure 1C**, RFP), indicating the successful induction of structural plasticity. After sLTP, Rab4^+^ endosomes lose processive long-range motion (**Figure 1C**, Rab4; **Video 2**). Using single-particle tracking (SPT; Sergé et al., 2008) to reconstruct motion trajectories and hidden Markov modeling with Bayesian model selection (HMM-Bayes; Monnier et al., 2015) to characterize motion types, we find that Rab4^+^ endosomes lose active transport, most prominently near the bases of sLTP-stimulated spines (**Figure 1C**, Rab4 and **Figure 1D**, Rab4; **Figure S2**, Rab4). sLTP triggers a similar reduction in active transport for Rab7^+^ and Rab11^+^ endosomes, and LAMP1^+^ lysosomes near the bases of targeted spines (**Figure 1D**; **Figure S2D**; **Figure S3A**; **Videos 3-5**). By contrast, ELs do not lose active transport at targeted spines in response to mock stimulation (sLTP stimulation in the absence of MNI; **Figure S3B-C**; **Video 6**), demonstrating that the induction of plasticity, not the application of laser, alters the motion of ELs. Glycine-induced chemical long-term potentiation (cLTP; Wong et al., 2024) also leads to the loss of active transport (**Figure S4**, cLTP versus Control), confirming the finding that stimulating plasticity inhibits the motion of ELs. Importantly, sLTP results in a higher number of ELs near the bases of stimulated spines (**Figure 1E**; **Figure S5** for antibody labeling), suggesting that blocking active transport concentrates ELs near spines undergoing plasticity.

We next asked whether ELs pause only briefly (<1 minute), as a transient byproduct of plasticity induction, or remain stationary for longer periods (>30 minutes), consistent with stable positioning that supports plasticity. The number of ELs near the bases of stimulated spines and the motion types of ELs do not change significantly even 30 minutes after sLTP (**Figure S6A-B**), suggesting long-term retention. Because technical limitations prevent us from tracking the same ELs over long periods, we cannot exclude the possibility that ELs paused immediately after sLTP are later replaced by different ELs (see **Methods**). Consequently, we used HMM-Bayes to determine the motion state-transition probabilities for ELs (**Figure S6C-D**; Monnier et al., 2015) and find that paused ELs are significantly less likely to resume active transport after sLTP (**Figure S6E**), indicating that ELs are not transported away from the bases of spines within 30 minutes of stimulation. Thus, sLTP not only inhibits EL transport but also concentrates and retains ELs near the bases of stimulated spines, supporting the hypothesis that ELs are stably positioned to support plasticity.

### Actin polymerization in the dendritic shaft is required to position endosomes and lysosomes near the bases of stimulated spines

We previously demonstrated that F-actin in the dendritic shaft confines AMPAR-containing vesicles to promote the exocytosis of AMPARs near sLTP-stimulated spines (Wong et al., 2024). We also found that actin polymerization inhibits the motion of rheological probes in the dendritic shaft (Wong et al., 2024), leading us to hypothesize that F-actin might have a general role in concentrating important proteins and organelles near spines to support plasticity. We therefore tested whether actin polymerization in the dendritic shaft also positioned ELs near sLTP-stimulated spines.

To visualize actin polymerization, we use F-tractin tagged with a fluorescent protein (Tractin) as a marker for F-actin (Schell et al., 2001). sLTP increases the fluorescence intensity of Tractin in spine heads and in dendritic shafts near the bases of targeted spines by stimulating actin polymerization in these regions (**Figure 2A**; **Figure S7**; Wong et al., 2024). We analyzed the motion of ELs inside areas with increased Tractin after sLTP (**Figure 2B**; **Figure S8**; **Videos 7-10**) and observed an increase in the number of ELs inside areas where sLTP stimulated the expansion of Tractin when compared to the same area prior to sLTP (**Figure 2C**, top row, black lines). By contrast, we do not observe an increase in the number of ELs outside regions with actin polymerization (**Figure 2C**, bottom row, black lines). Importantly, ELs inside areas with Tractin expansion lose active transport (**Figure 2D**, black lines), indicating the number of ELs increases inside areas with actin polymerization because they are retained (i.e., not transported away) and not because more ELs are transported through the imaged region.

**Figure 2.**
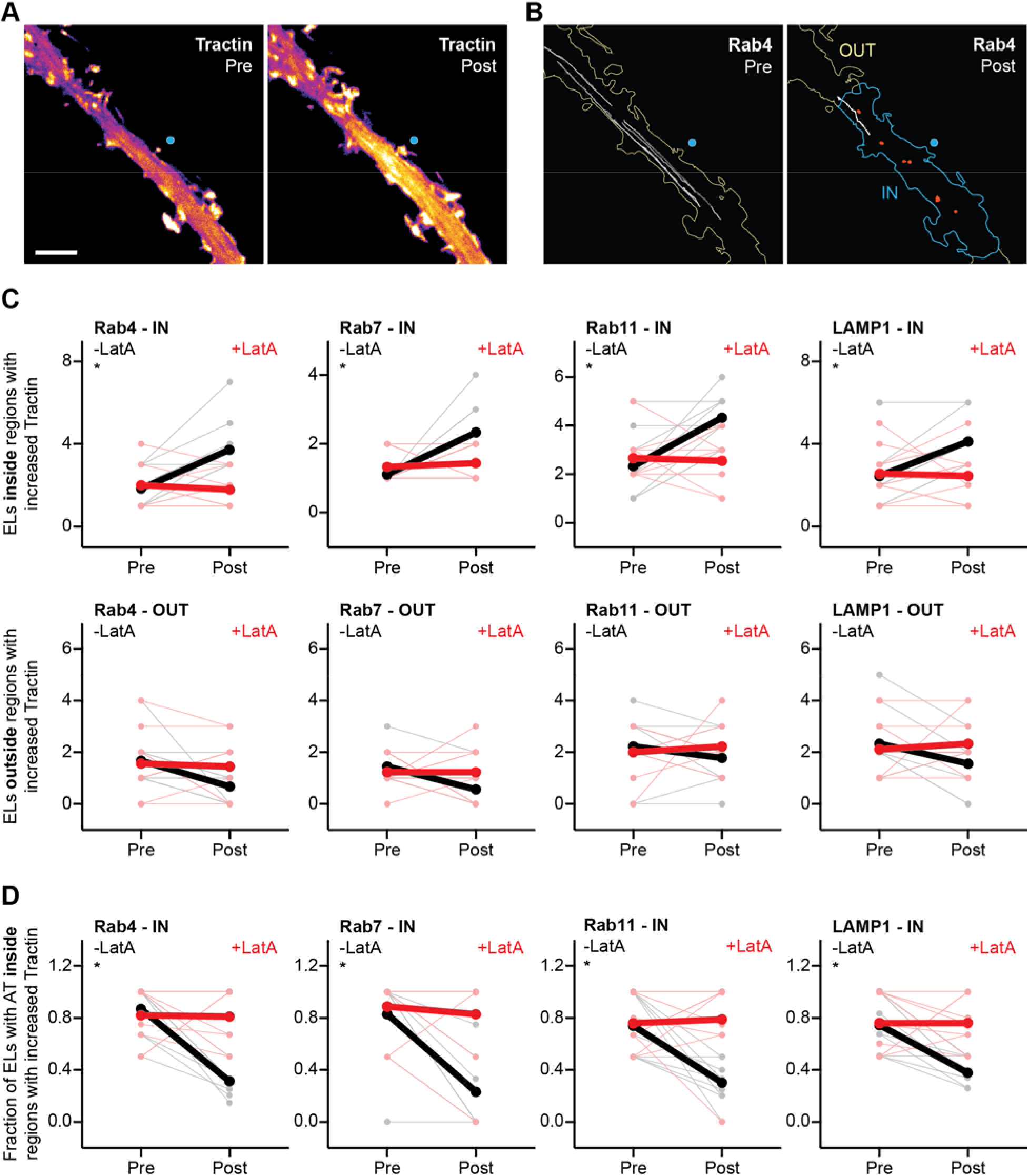
Actin polymerization in the dendritic shaft is required to position endosomes and lysosomes near the bases of sLTP-stimulated spines. (A-B) Representative spinning disk confocal images of F-tractin-mScarlet (Tractin; A) and Rab4-mEmerald^+^ (Rab4^+^) endosome trajectories (B) in the same dendritic shaft before and after structural long-term potentiation (sLTP) stimulation (Pre and Post, respectively). Scale bar, 5 µm. Blue dot indicates MNI uncaging site. Blue outline indicates region with increased Tractin signal. Gray trajectories indicate active transport; red trajectories indicate diffusive motion. (C) Top row: number of mEmerald-tagged Rab4^+^, Rab7^+^, or Rab11^+^ endosomes, or LAMP1^+^ lysosomes (ELs) inside regions with actin polymerization pre- and post-sLTP. Each pair of light gray dots represents the number of ELs in a dendrite pre- and post-sLTP (-LatA) for one stimulation experiment. Light red dots represent the number of ELs pre- and post-sLTP in the presence of latrunculin A (+LatA). Thick black and red dots represent the mean number of ELs pre- and post-sLTP in the absence or presence of latrunculin A, respectively. Rab4-IN -LatA, *p=0.0078; Rab7-IN -LatA, *p=0.0156; Rab11-IN -LatA, *p=0.0156; LAMP1-IN -LatA, *p=0.0078. Rab4-IN +LatA, p=0.6875; Rab7-IN +LatA, p>0.9999; Rab11-IN +LatA, p=0.9062; LAMP1-IN +LatA, p>0.9999. Significance determined by Wilcoxon signed-rank test with Holm–Šídák correction across four EL markers. Interaction, ΔEL# (Post – Pre), -LatA vs +LatA: Rab4-IN, **p=0.0006; Rab7-IN, *p=0.0341; Rab11-IN, *p=0.0081; LAMP1-IN, *p=0.0095 (see **Methods**). Significance determined by Mann-Whitney test with Holm–Šídák correction across four EL markers. Bottom row: number of ELs outside regions with actin polymerization pre- and post-sLTP. Rab4-OUT -LatA, p=0.0312; Rab7-OUT -LatA, p=0.0312; Rab11-OUT -LatA, p=0.3594; LAMP1-OUT -LatA, p=0.1094. Rab4-OUT +LatA, p=0.6719; Rab7-OUT +LatA, p>0.9999; Rab11-OUT +LatA, p>0.9999; LAMP1-OUT +LatA, p=0.7500. Significance determined by Wilcoxon signed-rank test with Holm–Šídák correction across four EL markers. Interaction, ΔEL# (Post – Pre), -LatA vs +LatA: Rab4-OUT, p=0.1180; Rab7-OUT, p=0.1636; Rab11-OUT, p=0.4696; LAMP1-OUT, p=0.1835 (see **Methods**). Significance determined by Mann-Whitney test with Holm–Šídák correction across four EL markers. N = 18 sLTP experiments for each marker (9 for -LatA and 9 for +LatA). For each experiment, the motion of ELs in the dendrite was imaged before and after sLTP across the entire field of view. Trajectories were then binned into regions with or without increased actin polymerization. Because we do not observe the expansion of Tractin after sLTP in the presence of latrunculin A, we analyzed the 20 µm region of dendritic shaft surrounding the stimulated spine (see **Figure S7C**). A new neuron was selected for each sLTP stimulation experiment; a maximum of three spines (i.e., neurons) was stimulated per neuronal culture. (D) Fraction of ELs exhibiting active transport (AT) inside regions with Tractin expansion pre- and post-sLTP. Each pair of light gray dots represents the fraction of ELs exhibiting AT inside a region with Tractin expansion in a dendrite pre- and post-sLTP (-LatA) for one stimulation experiment. Light red dots represent the fraction of ELs exhibiting AT pre- and post-sLTP in the presence of latrunculin A (+LatA). Thick black and red dots represent the mean fraction of ELs exhibiting AT pre- and post-sLTP in the absence or presence of latrunculin A, respectively. Rab4-IN -LatA, *p=0.0039; Rab7-IN -LatA, *p=0.0078; Rab11-IN -LatA, *p=0.0078; LAMP1-IN - LatA, *p=0.0078. Rab4-IN +LatA, p>0.9999; Rab7-IN +LatA, p>0.9999; Rab11-IN +LatA, p=0.6562; LAMP1-IN +LatA, p=0.9531. Significance determined by Wilcoxon signed-rank test with Holm–Šídák correction across four EL markers. Interaction, ΔEL AT (Post – Pre), -LatA vs +LatA: Rab4-IN, **p=0.0005; Rab7-IN, *p=0.0050; Rab11-IN, *p=0.0072; LAMP1-IN, *p=0.0213 (see **Methods**). Significance determined by Mann-Whitney test with Holm–Šídák correction across four EL markers. N = 18 sLTP experiments for each marker (9 for -LatA and 9 for +LatA). For each experiment, the motion of ELs in the dendrite was imaged before and after sLTP across an entire field of view. The fraction of trajectories exhibiting active transport inside regions with increased actin polymerization was then determined. Because we do not observe the expansion of Tractin after sLTP in the presence of latrunculin A, we analyzed the 20 µm region of dendritic shaft surrounding the stimulated spine (see **Figure S7C**). A new neuron was selected for each sLTP stimulation experiment; a maximum of three spines (i.e., neurons) was stimulated per neuronal culture.

Next, we tested whether EL positioning requires actin polymerization by examining the motion of ELs in neurons treated with latrunculin A (LatA), an actin depolymerization agent. Blocking actin polymerization with latrunculin A inhibits the accumulation of ELs near stimulated spines (i.e., the area where Tractin expansion would occur in the absence of latrunculin A; **Figure 2C**, top row, red lines) but has no significant effect on the number of ELs distal to stimulated spines (**Figure 2C**, bottom row, red lines). In addition, latrunculin A treatment abrogates the inhibitory effect of both sLTP and cLTP on the active transport of ELs (**Figure 2D**, red lines, for sLTP; **Figure S4**, cLTP+LatA for cLTP). Together, these results demonstrate that actin polymerization is required for the accumulation and retention of ELs near sLTP-stimulated spines.

Lastly, we tested whether actin polymerization itself, in the absence of other signaling pathways activated during plasticity, could inhibit the active transport of ELs. When neurons are treated with jasplakinolide (Jasp), a compound that induces actin polymerization and stabilizes F-actin, we find a significant reduction in the fraction of ELs exhibiting active transport (**Figure S4**, Jasp). Combined, these results demonstrate that sLTP induces actin polymerization in the dendritic shaft, which in turn positions ELs near stimulated spines.

### F-actin-dependent positioning of endosomes and lysosomes near the bases of stimulated spines predicts the magnitude of AMPAR turnover

Having demonstrated that actin polymerization positions components of the endocytic pathway near sLTP-stimulated spines, we asked whether the localization of ELs near target spines is required for plasticity or correlates with molecular and structural changes to spines after stimulation. Changes to AMPAR concentrations inside spines underlie the sustained increases or decreases in excitatory postsynaptic currents that characterize functional plasticity (Anggono and Huganir, 2012; Diering and Huganir, 2018). Consequently, we measured the turnover of GluA1 – an AMPAR subunit – in sLTP-stimulated spines and compared it with EL localization.

To quantify GluA1 turnover, we tagged GluA1 expressed from its endogenous locus with HaloTag (GluA1-HT) using homology-independent targeted integration (HITI; Suzuki et al., 2016; Wong et al., 2024). We then utilized a pulse-chase protocol with cell-impermeable Janelia Fluor (JF) dyes (Xie et al., 2017; Jonker et al., 2020) to label GluA1-HT on the surface of spines before and after sLTP (**Figure 3A**; **Figure S9**). By comparing the concentrations of GluA1-HT on the surface of a spine before sLTP (pulse; JF_635_i) and after sLTP (chase; JF_549_i), we can estimate the GluA1-HT turnover (defined here as the ratio of JF_549_i to JF_635_i) and correlate turnover with the localization of ELs (**Figure 3B**). The absence of ELs near the base of a spine before stimulation does not prevent initial spine enlargement in response to sLTP (**Figure S10A** and **S10B**, sLTP, left), but EL positioning after sLTP is correlated with the maintenance of spine volume over time (**Figure S10B**, sLTP +30 min, right). This finding suggests that the recruitment of ELs is not required for the induction phase of plasticity but instead supports the maintenance of plasticity (i.e., the post-enlargement phase during which structural and functional changes are stabilized). We can therefore target spines at which ELs are not present prior to stimulation to determine whether the recruitment of ELs in response to the induction of plasticity correlates with GluA1-HT turnover (**Figure 3B**, Rab4, Pre vs Post).

**Figure 3.**
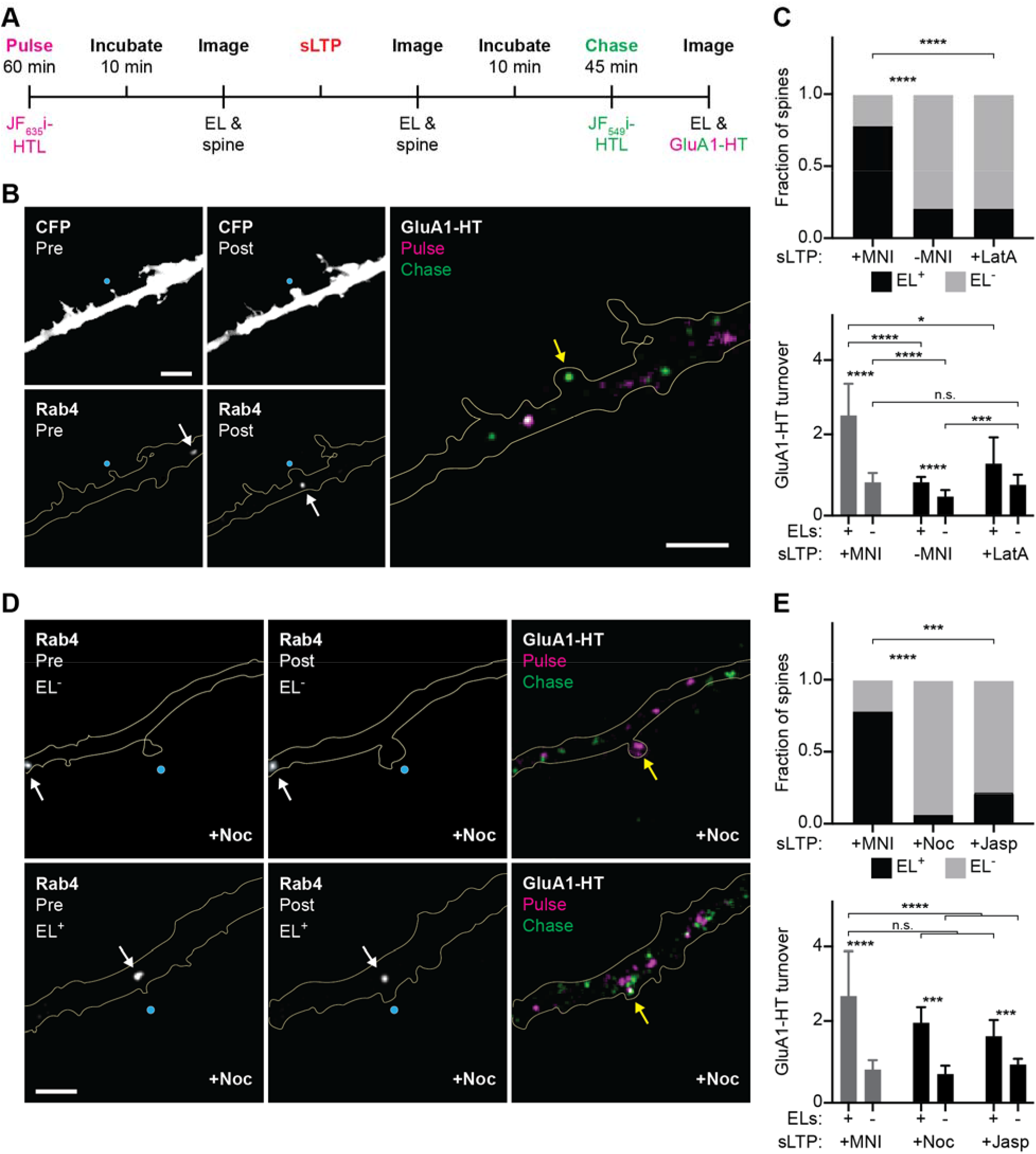
Actin-dependent positioning of endosomes and lysosomes near the bases of sLTP-stimulated spines predicts turnover of AMPA receptors. (A) Pulse-chase experiment to determine the localization of mEmerald-tagged Rab4^+^, Rab7^+^, or Rab11^+^ endosomes, or LAMP1^+^ lysosomes (ELs) and estimate the turnover of GluA1-HaloTag (GluA1-HT) in dendritic spines. GluA1-HT on the cell membrane is first labeled with the cell-impermeable JF_635_i dye conjugated to HaloTag ligand (JF_635_i-HTL). A target spine with no ELs near its base is then identified. Structural long-term potentiation (sLTP) stimulation is applied to the target spine and both the spine and EL markers are imaged to ensure successful induction of plasticity. Newly exocytosed GluA1-HT is then labeled with cell-impermeable JF_549_i-HTL. Finally, ELs and labeled GluA1-HT are imaged (see **Methods**). (B) CFP (left top): representative widefield images of a dendritic spine expressing mTurquoise2 immediately pre-sLTP (Pre) and post-sLTP (Post). Blue dot indicates MNI uncaging site. mTurquoise2 was used to identify and segment the dendritic spine and shaft. Rab4 (left bottom): widefield images of Rab4-mEmerald^+^ (Rab4^+^) endosomes inside the dendrite depicted above (CFP, left top) pre- and post-sLTP. White arrows indicate Rab4^+^ endosomes. GluA1-HT (right): widefield image of GluA1-HT on the surface of the dendrite depicted left (CFP, left top) labeled with JF_635_i-HTL before sLTP (Pulse, magenta) and then labeled with JF_549_i-HTL dye after sLTP (Chase, green). Yellow arrow indicates GluA1-HT on the sLTP-stimulated spine. Scale bars, 5 µm. (C) Top bar graph: fraction of spines that have ELs near their bases (EL^+^, black bar) after sLTP (+MNI), mock sLTP in the absence of MNI (-MNI), or sLTP in the presence of latrunculin A (+LatA). +MNI vs -MNI, ****p<0.0001; +MNI vs +LatA, ****p<0.0001. Significance determined by Fisher’s exact test with Holm–Šídák correction across the comparisons shown in graph. Bottom bar graph: turnover for GluA1-HT (JF_549_i-HTL/JF_635_i-HTL) on the surfaces of stimulated spines depending on whether ELs are positioned at their bases (EL: +/-) after sLTP. Bars and error bars represent mean and standard deviation, respectively. +MNI: EL^+^ vs EL^-^, ****p<0.0001; -MNI: EL^+^ vs EL^-^, ****p<0.0001; +MNI EL^-^ vs -MNI EL^-^, ****p<0.0001; +MNI EL^+^ vs -MNI EL^+^, ****p<0.0001; +MNI EL^+^ vs +LatA EL^+^, ***p=0.0002; +MNI EL^-^ vs +LatA EL^-^, p=0.5186; -MNI EL^-^ vs +LatA EL^-^, *p=0.0078. n.s., not significant. Significance determined by Mann-Whitney test with Holm–Šídák correction across the comparisons shown in graph. N = 18–60 sLTP-stimulated spines for each condition pooled from 4–16 sLTP experiments each for Rab4, Rab7, Rab11, or LAMP1 (see **Figure S11A**). For pulse-chase experiments analyzed in (C), we applied stimulation to dendritic spines that did not have ELs near their bases prior to stimulation. For each experiment, images of the spine and EL marker were taken immediately before and after sLTP stimulation, while images of the EL marker and pulse-chase labeling were taken after the chase (see **Methods**). A new neuron was selected for each sLTP stimulation experiment; a maximum of two spines (i.e., neurons) was stimulated per neuronal culture. EL^+^ vs EL^-^ turnover under +MNI and -MNI was additionally analyzed with a linear mixed-effects model to confirm results were not driven by non-independence among spines (+MNI, ****p<0.0001; -MNI, ****p<0.0001; see **Methods**). (D) Rab4 (left): representative widefield images of Rab4^+^ endosomes pre- and post-sLTP in the presence of nocodazole (+Noc). Blue dot indicates MNI uncaging site. White arrows indicate Rab4^+^ endosomes. Scale bar, 5 µm. GluA1-HT (right): widefield images of GluA1-HT on the surfaces of dendrites labeled with JF_635_i-HTL dye before sLTP (Pulse, magenta) and then labeled with JF_549_i-HTL dye after sLTP (Chase, green). Yellow arrow indicates GluA1-HT on the sLTP-stimulated spine. Scale bar, 5 µm. Nocodazole treatment inhibits EL positioning in response to sLTP. Top row depicts a dendritic spine without Rab4^+^ endosomes near its base both before and after sLTP (EL^-^) in the presence of nocodazole. Bottom row depicts a dendritic spine with a Rab4^+^ endosome near its base both before and after sLTP (EL^+^) in the presence of nocodazole. (E) Top bar graph: fraction of spines that have ELs near their bases (EL^+^, black bar) after sLTP in the presence of nocodazole (+Noc) or jasplakinolide (+Jasp). +MNI vs +Noc, ****p<0.0001; +MNI vs +Jasp, ***p=0.0001. Significance determined by Fisher’s exact test with Holm–Šídák correction across the comparisons shown in graph. +MNI data is reproduced from Figure 3C, top, +MNI. Bottom bar graph: turnover for GluA1-HT on the surfaces of stimulated spines depending on whether ELs are positioned at their bases before and after sLTP. Bars and error bars represent mean and standard deviation, respectively. +MNI EL^+^ vs +MNI EL^-^, ****p<0.0001; +Noc: EL^+^ vs EL^-^, ****p=0.0001; +Jasp: EL^+^ vs EL^-^, ***p=0.0001; +MNI EL^+^ vs +Noc EL^+^, p=0.1422; +MNI EL^+^ vs +Jasp EL^+^, p=0.0620. +MNI EL^+^ vs +Noc EL^-^, ****p<0.0001; +MNI EL^+^ vs +Jasp EL^-^, ****p<0.0001. n.s., not significant. Significance determined by Mann-Whitney test with Holm–Šídák correction across the comparisons shown in graph. N = 19–25 sLTP-stimulated spines for each condition pooled from 3–4 experiments each for Rab4, Rab7, Rab11, or LAMP1. For all pulse-chase experiments analyzed in (E, bottom), EL positioning did not change in response to sLTP. For EL^+^ experiments, we identified spines with ELs near target bases prior to stimulation. +MNI EL^+^ is a separate data set from Figure 3C, bottom. For EL^-^ experiments, we identified spines without ELs near target bases prior to stimulation. +MNI EL^-^ data is reproduced from Figure 3C, bottom. For each experiment, images of the spine and EL marker were taken immediately before and after sLTP stimulation, while images of the EL marker and pulse-chase labeling were taken after the chase (see **Methods**). For nocodazole and jasplakinolide treatments, a new neuronal culture was used for each experiment.

sLTP results in EL positioning near the bases of most, but not all, targeted spines (**Figure 3C**, top, +MNI), enabling us to compare GluA1-HT turnover in spines with versus without EL positioning post stimulation. Spines with ELs near their bases in response to sLTP had significantly greater GluA1-HT turnover than spines without ELs (**Figure 3C**, bottom, +MNI; **Figure S11A**, +MNI), demonstrating that EL positioning correlates with GluA1-HT turnover.

Mock sLTP stimulation (-MNI) fails to position ELs at the bases of most spines (**Figure 3C**, top, -MNI). Nevertheless, GluA1-HT turnover in spines with ELs positioned near their bases is modestly higher than in spines without ELs after mock sLTP stimulation (**Figure 3C**, bottom, - MNI; **Figure S11A**, -MNI), suggesting that ELs also support basal GluA1-HT turnover. Notably, sLTP results in a small but significant increase in GluA1-HT turnover even if ELs are not positioned in response to stimulation (**Figure 3C**, bottom, +MNI EL^-^ vs -MNI EL^-^). This increase in GluA1-HT turnover may be due to increased synaptic capture of newly exocytosed GluA1-HT diffusing into the stimulated spine from an adjacent region on the dendrite. However, the fold-increase in GluA1-HT turnover after sLTP is significantly greater in spines with EL positioning than in spines without EL positioning (i.e., turnover in sLTP-stimulated spines relative to mock-stimulated spines with the same EL status; **Figure S11B**). Combined, these findings reinforce our hypothesis that EL positioning is not required for the induction of plasticity but also demonstrate that EL positioning is strongly correlated with large increases in GluA1-HT turnover after the induction of plasticity.

Nevertheless, this correlation may reflect properties that differ between spines and influence both EL positioning and GluA1-HT turnover. We therefore asked whether this association holds when EL positioning is blocked pharmacologically at spines during sLTP. Latrunculin A treatment disrupts EL positioning (**Figure 3C**, top, +LatA) but also inhibits spine enlargement (**Figure S12A-B**) and the large increase in GluA1-HT turnover associated with EL positioning after sLTP (**Figure 3C**, bottom, +LatA EL^+^ vs +MNI EL^+^; see **Figure S12C**, -MNI +LatA, for mock stimulation controls), raising the possibility that latrunculin A completely blocks the induction of plasticity in addition to actin-mediated EL positioning. However, latrunculin A does not reduce GluA1-HT turnover at sLTP-stimulated spines without EL positioning (**Figure 3C**, bottom, +LatA EL^-^ vs +MNI EL^-^ and +LatA EL^-^ vs -MNI EL^-^), indicating that treated spines respond to plasticity-inducing stimuli even though structural plasticity is absent. This observation is consistent with previous findings that inhibiting actin polymerization spares the induction of functional plasticity but impairs its maintenance (Krucker et al., 2000; Fukazawa et al., 2003), leading us to hypothesize that latrunculin A reduces GluA1-HT turnover by inhibiting actin-dependent processes that support plasticity after induction, including EL positioning, rather than by preventing induction itself. Because latrunculin A disrupts both EL positioning and structural plasticity, however, it cannot distinguish their contributions to GluA1-HT turnover. We therefore sought to identify alternative treatments to manipulate EL positioning without abolishing actin-dependent structural plasticity and then test whether these treatments would alter GluA1-HT turnover.

We previously demonstrated that nocodazole (Noc), a microtubule depolymerization agent, prevents long-ranged transport of AMPAR-containing vesicles (Wong et al., 2024). We reasoned that nocodazole treatment could be used to block microtubule-based transport of ELs to a stimulated spine and thus inhibit EL positioning in response to sLTP, unless ELs are present at the base of the spine before stimulation. Indeed, nocodazole disrupts EL positioning near the bases of spines in response to sLTP if ELs are not present prior to stimulation (**Figure 3D**, top and **Figure 3E**, top, +Noc). Moreover, ELs near the bases of nocodazole-treated spines remain at their positions after sLTP (**Figure 3D**, bottom; **Figure S13A-B**). Importantly, nocodazole does not prevent structural plasticity in response to sLTP regardless of whether ELs are present (**Figure S13C**). Nocodazole significantly reduces GluA1-HT turnover after sLTP among spines lacking ELs prior to stimulation compared to untreated spines (**Figure S14**), leading us to ask whether this reduction was the result of inhibiting EL positioning. GluA1-HT turnover in nocodazole-treated spines without EL positioning before and after sLTP is reduced compared to turnover in untreated spines where ELs are positioned before and after sLTP (**Figure 3E**, bottom,

+MNI EL^+^ vs +Noc EL^-^; see **Figure S13D**, -MNI +Noc, for mock stimulation controls). By contrast, GluA1-HT turnover in nocodazole-treated spines with ELs present near their bases before, and thus after, sLTP is greater than turnover in spines without EL positioning (**Figure 3D** and **Figure 3E**, bottom, +Noc EL^+^ vs +Noc EL^-^). These results demonstrate that increased GluA1-HT turnover in response to sLTP is inhibited by nocodazole only under conditions where nocodazole has prevented EL positioning (i.e., if ELs are not present prior to stimulation). These findings can be recapitulated by treating neurons with jasplakinolide, which inhibits the transport of ELs to stimulated spines by promoting actin polymerization in the dendritic shaft (**Figure 3E**, top, +Jasp; **Figure S4**, Jasp) – rather than by disrupting microtubules – but also does not abolish structural plasticity (**Figure S15A-C**). Like nocodazole, jasplakinolide inhibits increased GluA1-HT turnover in response to sLTP only if ELs are absent from the bases of targeted spines before, and thus after, stimulation (**Figure 3E**, bottom, +MNI EL^+^ vs +Jasp EL^-^; +Jasp EL^+^ vs +Jasp EL^-^; **Figure S14**; see **Figure S15D**, -MNI +Jasp, for mock stimulation controls). These results show that preventing EL positioning rather than blocking structural plasticity inhibits the large increase in GluA1-HT turnover we observe in response to sLTP.

Overall, these findings demonstrate that GluA1-HT turnover is strongly associated with EL positioning even when positioning is manipulated with pharmacological agents. The correlation between GluA1-HT turnover and EL positioning is also observed when plasticity is induced with cLTP rather than sLTP (**Figure S16**). In summary, we find that the induction of plasticity stimulates actin polymerization in the dendritic shaft, which positions ELs near stimulated sites. Although EL positioning is not required for the induction of plasticity, it predicts a large increase in GluA1-HT turnover after the induction of plasticity, suggesting that the recruitment of ELs supports the maintenance of plasticity.

## Discussion

In this study, we use a combination of photostimulation and single-particle tracking (SPT with HMM-Bayes; Sergé et al., 2008; Monnier et al., 2015) to examine the motion of the endosomes and lysosomes (ELs) in response to the induction of plasticity. We find that ELs lose motion, and are retained, near the bases of spines stimulated with a structural long-term potentiation (sLTP) protocol in a manner that is dependent on actin polymerization in the dendritic shaft. EL positioning is a strong predictor of the magnitude of α-amino-3-hydroxy-5-methyl-4-isoxazolepropionic acid receptor (AMPAR) turnover in sLTP-stimulated spines.

Together with previous work demonstrating critical roles for the endocytic pathway in regulating synaptic plasticity and strength (Ehlers, 2000; Park et al., 2004; Park et al., 2006; Brown et al., 2007; Kelly et al., 2011; Kneussel and Hausrat, 2016; Padamsey et al., 2017), these results are consistent with a model in which F-actin in the dendritic shaft supports plasticity by recruiting the endocytic machinery to the bases of stimulated spines.

These findings raise further questions regarding the precise mechanisms through which F-actin inhibits active transport. While ELs are transported along the length of dendrites via microtubule-based transport (Goo et al., 2017; Yap et al., 2018; Yap and Winckler, 2022), they are also transported short distances on F-actin via interactions with myosin (Correia et al., 2008; Wang et al., 2008; Esteves da Silva et al., 2015). Interactions with myosin or other proteins may anchor ELs to F-actin near spines in response to synaptic activity (Langford, 2002; van Bommel et al., 2019). Alternatively, F-actin may block the active transport of ELs indirectly. We and others have previously found actin polymerization inhibits motion inside dendritic shafts and spines by changing the rheological properties of the dendritic cytoplasm (Obashi et al., 2019; Wong et al., 2024). Consequently, F-actin can indirectly inhibit the active transport of AMPAR-containing vesicles (Wong et al., 2024).

The observation that F-actin generally inhibits motion along the length of the dendritic shaft indicates that it may act as an organizational hub for other organelles and macromolecular complexes that play roles in plasticity. Future studies should examine the distribution of other organelles (e.g., mitochondria) and proteins (e.g., CaMKII⍰) relative to F-actin in the dendritic shaft and evaluate how F-actin-mediated positioning contributes to plasticity. While our work finds a strong association between the presence of ELs and AMPAR turnover, it does not demonstrate definitively that EL positioning is required for increased AMPAR turnover.

Optogenetic tools that control the subcellular positioning of organelles and proteins (e.g., iLID and eMags; van Bergeijk et al., 2015; Benedetti et al., 2018; Benedetti et al., 2020; Nijenhuis et al., 2020) could be used to more precisely determine which molecules must be recruited to the bases of stimulated spines to support plasticity.

If F-actin inhibits motion in general, however, it should also impede cargo destined for other spines. A dendrite contains numerous spines, each displaying a different level of activity and supported by a unique complement of proteins and organelles (Berry and Nedivi, 2017). If all passing cargo were trapped at a single stimulated spine, neurons could not support plasticity at multiple synapses independently. Mechanisms must therefore exist that allow unrelated cargo to pass unhindered. One possibility is that the assortment of motors decorating a cargo determines whether it is retained by F-actin or passes through (Ferro et al., 2019). Defining these mechanisms would improve our understanding of how neurons maintain input specificity.

Synaptic protein recycling via the endocytic pathway must be carefully regulated to maintain an adequate reservoir of synaptic proteins to support neuronal activity. Here we find evidence that the function of the endocytic pathway is also regulated by its dynamic localization. Continuing to elucidate the precise localization of the endocytic pathway, and the timeline by which components of this pathway are assembled, in response to stimulation will enhance our understanding of synaptic plasticity and thus learning and memory.

## Materials and methods

Animal work was conducted according to the Institutional Animal Care and Use Committee guidelines of Janelia Research Campus (IACUC protocol #21-0206).

### Key resources table

| Reagent type (species) or resource | Designation | Source or reference | Identifiers | Additional information |
| --- | --- | --- | --- | --- |
| Antibody | Chicken anti-GFP (anti-mEmerald) | Aves Labs | GFP-1010 | IF(1:2000)<br>Reacts with mEmerald |
| Antibody | Goat anti-chicken STAR GREEN | abberior instruments GmbH | STGREEN-1005-500UG | IF(1:500) |
| Antibody | Goat anti-mouse ATTO 647N | Rockland Immunochemical, Inc. | 610-156-121 | IF(1:500) |
| Antibody | Goat anti-rabbit ATTO 594 | Rockland Immunochemical, Inc. | 611-155-122 | IF(1:500) |
| Antibody | Mouse anti-EEA1 | Thermo Fisher Scientific | 14-9114-82 | IF(1:500) |
| Antibody | Mouse anti-LAMP1 | Thermo Fisher Scientific | MA1-164 | IF(1:50) |
| Antibody | Mouse anti-Rab9 | Thermo Fisher Scientific | MA3-067 | IF(1:200) |
| Antibody | Mouse anti-Rab11A | Thermo Fisher Scientific | MA5-49197 | IF(1:100) |
| Antibody | Rabbit anti-EHD1 | Thermo Fisher Scientific | MA565176 | IF(1:50) |
| Antibody | Rabbit anti-LAMP2 | Thermo Fisher Scientific | PA1-655 | IF(1:50) |
| Antibody | Rabbit anti-Rab4 | Abcam | ab13252 | IF(1:200) |
| Antibody | Rabbit anti-Rab7A | Proteintech | 55469-1-AP | IF(1:50) |
| Chemical compound, drug | (-)-Bicuculline methiodide (bicuculline) | Tocris | 2503 | 20 μM |
| Chemical compound, drug | Bovine serum albumin (BSA) | Sigma-Aldrich | B4287-5G | 3% (w/v) |
| Chemical compound, drug | 1,4-diazabicyclo[2.2.2]octane (DABCO) | Sigma-Aldrich | D27802 | 2.5% (w/v) |
| Chemical compound, drug | Dimethyl sulfoxide (DMSO), anhydrous | Thermo Fisher Scientific | D12345 |  |
| Chemical compound, drug | Glycine | Tocris | 0219 | cLTP: 0.2 mM<br>IF: 0.1 M |
| Chemical compound, drug | Hibernate A – Low Fluorescence | Transnetyx | HALF500 |  |
| Chemical compound, drug | Jasplakinolide (Jasp) | Tocris | 2792 | 100 nM |
| Chemical compound, drug | JF<sub>549</sub>i-HTL | Xie et al., 2017 | N/A | 50 nM |
| Chemical compound, drug | JF<sub>635</sub>i-HTL | Jonker et al., 2020 | N/A | 50 nM |
| Chemical compound, drug | Latrunculin A (LatA) | Tocris | 3973 | 1 μM |
| Chemical compound, drug | MNI-caged-L-glutamate (MNI) | Tocris | 1490 | 2 mM |
| Chemical compound, drug | Mowiol 4-88 | Sigma-Aldrich | 813831 | 10% (w/v) |
| Chemical compound, drug | NbActiv4 Minus Phenol Red | Transnetyx | NB4PR500 |  |
| Chemical compound, drug | Nocodazole (Noc) | Tocris | 1228 | 5 μM |
| Chemical compound, drug | Normal goat serum | Thermo Fisher Scientific | 31873 | 5% (w/v) |
| Chemical compound, drug | 16% paraformaldehyde (PFA) | Electron Microscopy Sciences | 15713 | 4% |
| Chemical compound, drug | Poly-D-lysine (PDL) hydrobromide | Sigma-Aldrich | P6407 | 0.5 mg/mL |
| Chemical compound, drug | ProLong™ Gold | Thermo Fisher Scientific | P10144 |  |
| Chemical compound, drug | Silicon rhodamine (SiR)-actin | Cytoskeleton, Inc. | CY-SC001 | 500 nM |
| Chemical compound, drug | Strychnine hydrochloride | Sigma-Aldrich | S8753 | 1 μM |
| Chemical compound, drug | Tetrodotoxin citrate (TTX) | Tocris | 1069 | 1 μM |
| Chemical compound, drug | Triton X-100 | Sigma-Aldrich | T8787 | 0.1% |
| Chemical compound, drug | Tween-20 | Sigma-Aldrich | P9416 | 0.05% |
| Commercial assay or kit | P3 Primary Cell 96-well Nucleofector™ Kit (96 RCT) | Lonza | V4SP-3096 |  |
| Recombinant DNA reagent | pAAV-hSyn1-mTurquoise2 | Viviana Gradinaru | Addgene #99125 |  |
| Recombinant DNA reagent | pAAV-hSyn-eGFP-WPRE-bGH | James Wilson | Addgene #105539 |  |
| Recombinant DNA reagent | pAAV-hSyn-tdTomato | Erin O'Shea | Addgene #239048 |  |
| Recombinant DNA reagent | mEmerald-Rab4a-7 | Michael Davidson | Addgene #54242 |  |
| Recombinant DNA reagent | mEmerald-Rab7a-7 | Michael Davidson | Addgene #54244 |  |
| Recombinant DNA reagent | mEmerald-Rab11a-7 | Michael Davidson | Addgene #54245 |  |
| Recombinant DNA reagent | mEmerald-Lysosomes-20 (LAMP1) | Michael Davidson | Addgene #54149 |  |
| Recombinant DNA reagent | F-tractin-mScarlet-I | This paper | Addgene #263441 |  |
| Recombinant DNA reagent | PX551 | Swiech et al., 2015 | Addgene #60957 |  |
| Recombinant DNA reagent | px552-sg-Gria1-HT | Wong et al., 2024 | Addgene #187652 |  |
| Recombinant DNA reagent | hSyn-tractin | Wong et al., 2024 | Addgene #187649 |  |
| Software, algorithm | 3C/S2D/S11B/S16B analysis code | This paper | https://doi.org/10.5281/zenodo.23073463 |  |
| Software, algorithm | Claude (claude-opus-5-5) | Anthropic (San Francisco, CA) | https://claude.ai |  |
| Software, algorithm | Elements C Imaging Software, Version 5.21.00 | Nikon Instruments (Melville, NY) | N/A |  |
| Software, algorithm | Fiji | Schindelin et al., 2012 | N/A |  |
| Software, algorithm | HMM-Bayes | Monnier et al., 2015 | https://github.com/lcbb/HMM-Bayes | MATLAB script used to predict motion states of particles |
| Software, algorithm | Imaris 9.3 | Bitplane (Belfast, United Kingdom) | N/A |  |
| Software, algorithm | iMSPECTOR | abberior instruments GmbH (Göttingen, Germany) | N/A |  |
| Software, algorithm | MATLAB R2015b | MathWorks (Natick, MA) | https://www.mathworks.com/ |  |
| Software, algorithm | MSDAnalyzer | Tarantino et al., 2014 | https://github.com/tinevez/msd-analyzer | Packaged into SLIMfast |
| Software, algorithm | NumPy 2.4.4 | Harris et al., 2020 | https://numpy.org | Python package used for statistical analysis |
| Software, algorithm | Pandas 3.0.2 | McKinney, 2010 | https://pandas.pydata.org | Python package used for data handling |
| Software, algorithm | Patsy 1.0.3 |  | https://github.com/pydata/patsy | Python package used for statistical analysis |
| Software, algorithm | Prism 11 | GraphPad (San Diego, CA) | https://www.graphpad.com/ |  |
| Software, algorithm | Python 3.11.15 | Python Software Foundation | https://www.python.org |  |
| Software, algorithm | SciPy 1.17.1 | Virtanen et al., 2020 | https://scipy.org | Python package used for statistical analysis |
| Software, algorithm | SLIMfast | Wong et al., 2024 | https://doi.org/10.5281/zenodo.14901152 | 2D single-particle tracking MATLAB program |
| Software, algorithm | statsmodels 0.15.0 | Seabold and Perktold, 2010 | https://www.statsmodels.org | Python package used for statistical analysis |
| Strain, strain background (AAV) | rh10-hSyn1-mTurquoise2 | Janelia Research Campus Viral Services | N/A |  |
| Strain, strain background (AAV) | rh10-hSyn-tdTomato | Janelia Research Campus Viral Services | N/A |  |
| Strain, strain background (AAV) | rh10-hSyn-GFP | Janelia Research Campus Viral Services | N/A |  |
| Strain, strain background (*Rattus norvegicus*) | Sprague-Dawley rats, CD IGS (Sprague-Dawley postnatal pups P0, M and F) | Charles River Laboratories | Strain code: 400 |  |
| Strain, strain background<br>(*Escherichia coli*) | Stbl3 | Invitrogen | C737303 |  |

### Plasmids

mEmerald-tagged markers for endosomes and lysosomes (wild-type Rab4, Rab7, Rab11, and LAMP1) were gifts from Michael Davidson (obtained via Addgene). F-tractin-mScarlet was constructed by amplifying mScarlet-I and cloning it into the AgeI and BsrGI restriction sites of hSyn-tractin, thereby replacing mNeonGreen with mScarlet-I (Wong et al., 2024). Constructs used to insert HaloTag into *Gria1* (the gene encoding GluA1) via homology-independent targeted integration (HITI) were previously described (Swiech et al., 2015; Wong et al., 2024). All plasmids were propagated in Stbl3 *E. coli* cells.

### Neuronal cell culture and transfection

Dissociated hippocampal neurons were prepared from postnatal day 0 (P0) Sprague-Dawley rat pups (Charles River) and transfected via nucleofection as previously described (Nunez, 2008; Wong et al., 2024; Wong et al., 2025). Briefly, 500,000 hippocampal neurons were transfected with 0.5 µg of each target plasmid using an Amaxa 4D-Nucleofector with program CU-110 immediately after dissociation from P0 rat pups. After nucleofection, approximately 50,000–75,000 cells were spread onto the coverslips of poly-D-lysine (PDL)-coated 35 mm MatTek dishes or 35 mm Ibidi gridded µ-Dishes. Neurons were processed at 14–21 days in vitro (DIV 14–21).

### Homology-independent targeted integration (HITI)

HaloTag was inserted into the endogenous locus of *Gria1* via HITI as previously described (Wong et al., 2024; Wong et al., 2025). Briefly, 0.5 µg of px552-sg-Gria1-HT and 0.5 µg of PX551 were delivered into freshly dissected P0 rat hippocampal neurons via nucleofection prior to plating. Transfected neuronal cultures were used for labeling at 14–21 days in vitro (DIV 14–21). The expression, trafficking, and function of GluA1-HaloTag (GluA1-HT) were previously validated (Wong et al., 2024).

### Adeno-associated virus (AAV) transduction

hSyn-tdTomato, hSyn-GFP, and hSyn1-mTurquoise2 – used to express fluorescent cytosolic markers to define dendrites and dendritic spines – were packaged into adeno-associated virus (AAV) by the Janelia Research Campus Viral Services Shared Resource as previously described (Wong et al., 2024). Cultured rat hippocampal neurons were infected with 1×10^8^ genomic copies of rh10-hSyn-tdTomato, rh10-hSyn-GFP, or rh10-hSyn1-mTurquoise2 at 3–7 days in vitro (DIV 3–7).

### Microscopes

Photostimulation experiments and live-cell imaging for single-particle tracking (SPT) were performed on either a Nikon Eclipse TiE inverted microscope or a Nikon CSU-W1 inverted spinning disk confocal microscope. The Nikon Eclipse TiE inverted microscope is equipped with 405/445/488/514/561/594/642 nm laser lines for automatic total internal reflection fluorescence (TIRF) microscopy, a focused 405 nm laser for photomanipulation (∼5 mW at sample at 100% power; ∼1 µm diameter focal spot), three iXon Ultra 897 electron-multiplying charge-coupled device (EMCCD) cameras connected via a tri-cam splitter for simultaneous multicolor acquisition, and a TIRF illuminator. Highly inclined and laminated optical sheet (HILO) imaging was performed using the TIRF illuminator as previously described (Wong et al., 2024; Wong et al., 2025). The Nikon CSU-W1 inverted spinning disk confocal microscope (50 µm-pinholed disk) is equipped with 405/445/488/514/561/594/642 nm laser lines for confocal microscopy, 405/473 nm laser lines for galvo-based photomanipulation (∼6.7 mW at sample at 100% power; ∼1 µm diameter focal spot), and two Hamamatsu ORCA-Fusion BT scientific complementary metal-oxide-semiconductor (sCMOS) cameras. Each Nikon microscope system is equipped with a fast Z-piezo stage and Perfect Focus System. A Tokai Hit Stage Top Incubator was used to maintain constant environmental conditions of 5% CO_2_ and 37 °C during imaging for both Nikon microscope systems. For all experiments performed with these microscopes, we used a 100× Apochromat TIRF (APO TIRF) oil-immersion objective (numerical aperture [NA] = 1.49). Both Nikon microscope systems were controlled by Nikon Elements C Imaging Software.

Stimulated emission depletion (STED) imaging was performed on an abberior Expert Line STED microscope equipped with 405/440/485/561/640 nm laser lines for illumination and 595/775 nm laser lines for depletion. Emitted light was collected with four spectral avalanche photodiodes (APDs). A 60× Plan Apochromat oil-immersion objective (NA = 1.42) was used for all STED imaging. The STED microscope was controlled by abberior iMSPECTOR.

### Endosome and lysosome (EL) imaging

Prior to imaging, NbActiv4 neuronal cell culture medium was replaced with imaging buffer (150 mM NaCl, 2 mM CaCl_2_, 2 mM MgCl_2_, 5 mM KCl, 10 mM HEPES, 30 mM -glucose, pH 7.4). To track the motion of endosomes and lysosomes (ELs), and to avoid overexpression artifacts (Hornfeck et al., 2026), we selected neurons that expressed low-to-moderate levels of the mEmerald-tagged EL marker (i.e., neurons that had clearly defined fluorescent puncta and low levels of fluorescence background in the cytosol). Imaging particles using the Nikon Eclipse TiE inverted microscope was previously described (Wong et al., 2024; Wong et al., 2025). To image ELs using the Nikon CSU-W1 microscope, a dendrite of interest expressing an EL marker was first identified. A region of interest (ROI), typically 350 × 350 pixels to 512 × 512 pixels, was defined around the dendrite to enable faster imaging and to reduce file size. To ensure that ELs could be imaged as they traversed the length of the ROI, we selected dendrites with low variance in Z (i.e., dendrites that lie flat, parallel to the coverslip). The spinning disk speed was set to ultra-quiet mode. Finally, we adjusted the focal plane (z) so that the ELs were in focus and commenced imaging at 20 frames per second for 250–500 frames. We limited the number of frames imaged to reduce photobleaching and phototoxicity, as well as data storage requirements, especially for regions that required tracking over multiple time points (e.g., ELs that are tracked 30 minutes after sLTP stimulation).

### Photostimulation

The protocol for inducing structural long-term potentiation (sLTP) using 4-methoxy-7-nitroindolinyl-caged-L-glutamate (MNI) uncaging was previously validated and described (Wong et al., 2024). For all photostimulation experiments, neurons were washed three times with photostimulation buffer (150 mM NaCl, 2 mM CaCl_2_, 5 mM KCl, 10 mM HEPES, 30 mM - glucose, pH 7.4, plus 1 μM tetrodotoxin [TTX]). Photostimulation buffer was then replaced with 2 mM MNI in photostimulation buffer, and samples were placed onto either the Nikon Eclipse TiE or Nikon CSU-W1 microscope. Target spines were identified using a cytosolic fluorescent marker (e.g., tdTomato), and a 405 nm uncaging laser was parked ∼1 µm perpendicular to the tips of target spine heads. We selected healthy, well-isolated spines (>2 µm from their nearest neighboring spine; **Figure S2A-B**) to reduce the activation of neighboring spines (see Wong et al., 2024 for validation of single-spine stimulation). For the Nikon Eclipse TiE, the power of the 405 nm uncaging laser was set to 30% with a pulse width of 0.1 ms. For the Nikon CSU-W1, the power of the 405 nm uncaging laser was set to 5% with a pulse width of 0.1 ms. sLTP was induced by firing the 405 nm uncaging laser at 0.5 Hz for 120 seconds. Images of the target spine were taken before and after sLTP stimulation to confirm spine enlargement. For mock sLTP stimulation control experiments in the absence of MNI (-MNI), MNI was not added to photostimulation buffer prior to imaging, but optical parameters remained the same.

### Chemical treatments

The protocol for glycine-induced chemical long-term potentiation (cLTP) was previously described (Wong et al., 2024). Briefly, cultured neurons were washed three times with chemical stimulation buffer (150 mM NaCl, 2 mM CaCl_2_, 5 mM KCl, 10 mM HEPES, 30 mM -glucose, 20 µM bicuculline, 1 µM strychnine, pH 7.4) and then stimulated with 0.2 mM glycine in chemical stimulation buffer for 15 minutes at 37 °C and 5% CO_2_. Neurons were then rinsed twice with wash buffer (150 mM NaCl, 2 mM CaCl_2_, 2 mM MgCl_2_, 5 mM KCl, 10 mM HEPES, 30 mM -glucose, 20 µM bicuculline, 1 µM strychnine, pH 7.4) to remove glycine prior to imaging. To test the effect of F-actin stabilization on particle motion, neurons were treated with 100 nM jasplakinolide (Jasp) for 10 minutes prior to imaging or photostimulation. To examine the effect of actin depolymerization, neurons were treated with 1 µM latrunculin A (LatA) for 5 minutes prior to imaging or photostimulation. To test the effect of microtubule depolymerization on cargo transport, neurons were treated with 5 µM nocodazole (Noc) for 2 hours prior to imaging or photostimulation. For jasplakinolide-, latrunculin A-, and nocodazole-treatment experiments, the pharmacological agents remained in the culture during imaging or photostimulation.

### Immunofluorescence

To label endogenous EL markers and mEmerald, cultured neurons were fixed between 14–21 days in vitro (DIV 14–21). Neurons were fixed with 4% paraformaldehyde (PFA) in 1× phosphate-buffered saline (PBS) for 15 minutes. The fixative was quenched with 0.1 M glycine in 1× PBS for 10 minutes. Neurons were then washed three times for 5 minutes each wash with 1× PBS. The cell membranes of neurons were permeabilized with 0.1% Triton X-100 in 1× PBS for 5 minutes. After permeabilization, neurons were washed three times for 5 minutes each wash with 1× PBS and then blocked in blocking buffer (3% bovine serum albumin, 5% normal goat serum, and 0.05% Tween-20 in 1× PBS) for 60 minutes at room temperature. EL markers were labeled with primary antibodies (dilutions listed in **Key resources table**) in blocking buffer overnight at 4 °C. After primary antibody labeling, neurons were washed once in blocking buffer for 10 minutes and three times in PBST (0.05% Tween-20 in 1× PBS) for 5 minutes each wash. Neurons were then labeled with secondary antibodies (dilutions listed in **Key resources table**) in blocking buffer for 45 minutes at room temperature. Lastly, neurons were washed three times for 5 minutes each wash with 1× PBS and mounted in ProLong Gold antifade mountant.

Immunofluorescence imaging was performed on a Nikon CSU-W1 microscope using a 100× Apochromat TIRF oil-immersion objective. Z-stacks were recorded with a step size of 0.3 µm. The spinning disk speed was set to ultra-quiet mode. Experiments in which samples were fixed and labeled with antibodies after photostimulation required additional steps to ensure that targeted spines could be re-identified after labeling for immunofluorescence imaging. First, neurons were plated onto PDL-coated 35 mm Ibidi gridded µ-Dishes. Second, alignment markers were drawn on the sample dish and sample holder so that we could find the dish’s original orientation even after removing it from the sample holder. Lastly, we recorded the grid space designation for each targeted spine.

### F-actin imaging

To determine the spatial relationship between ELs and F-actin after sLTP, we stimulated neurons (see **Photostimulation**) expressing F-tractin-mScarlet and an EL marker. Like the cytosolic markers, F-tractin-mScarlet was imaged before and after sLTP to confirm actin polymerization and spine enlargement. To image F-actin by STED, we photostimulated neurons expressing F-tractin-mNeonGreen using the Nikon CSU-W1 (see **Photostimulation**), labeled F-actin with silicon rhodamine (SiR)-actin, and then imaged the stimulated spine and surrounding dendrite using the abberior Expert Line STED. To enable the re-identification of sLTP-stimulated spines after SiR-actin labeling, we used neurons plated onto PDL-coated 35 mm Ibidi gridded µ-Dishes. In addition, we used a sample holder that could be mounted onto the stages of both the Nikon CSU-W1 and the abberior Expert Line STED microscopes without adjustment. We drew alignment markers on the dish and sample holder so that we could return the dish to its original orientation after SiR-actin labeling. After stimulating a target spine and recording its grid space designation, we fixed the sample with 4% PFA in 1× PBS for 20 minutes. The fixative was quenched with 0.1 M glycine in 1× PBS for 10 minutes. Neurons were then washed three times for 5 minutes each wash with 1× PBS. Neurons were briefly permeabilized with 0.1% Triton X-100 in 1× PBS for 5 minutes and washed three times for 5 minutes each wash with 1× PBS. F-actin was then labeled using 500 nM SiR-actin for 60 minutes. Finally, SiR-actin was removed, and the sample was washed once with 1× PBS before being mounted using Mowiol 4-88 containing 1,4-diazabicyclo[2.2.2]octane (DABCO). After labeling F-actin, the sample was placed onto the abberior Expert Line STED microscope. Using a CoolLED illumination system and GFP filter sets to view F-tractin-mNeonGreen, target spines were re-identified based on recorded grid locations. ROIs were defined around each target spine, and confocal images of F-tractin-mNeonGreen for each ROI were taken to ensure the correct spines had been re-identified. Still using F-tractin-mNeonGreen, we defined the Z-range of the dendrite of interest. We then set up parameters for imaging SiR-actin. First, we selected the 640 nm laser line for illumination and the 775 nm donut laser for depletion. Next, we selected a detection range appropriate for SiR (655–700 nm). The pinhole size was set to 0.91 Airy units. A pixel size of 40 nm was used.

Finally, depletion laser power was set to 15% in xy and 5% in z. A confocal Z-stack of F-tractin-mNeonGreen was also recorded to define the boundaries of the dendrite.

### Pulse-chase labeling with Janelia Fluor (JF) dyes

Cell-impermeable Janelia Fluor 635 HaloTag ligand (JF_635_i-HTL) and cell-impermeable Janelia Fluor 549 HaloTag ligand (JF_549_i-HTL) were kind gifts from Dr. Luke Lavis. Dyes were reconstituted in DMSO at a concentration of 200 µM and diluted to working concentrations of 50 nM in appropriate stimulation or imaging buffers before use. Neurons expressing GluA1-HT were used for pulse-chase experiments at 14–21 days in vitro (DIV 14–21). First, NbActiv4 neuronal culture medium was removed and replaced with 50 nM JF_635_i-HTL in imaging buffer (pulse). Neurons were brought to the Nikon Eclipse TiE or Nikon CSU-W1 and labeled with JF_635_i-HTL in the stage-top incubators at 37 °C and 5% CO_2_ for 60 minutes. During this time, target spines were identified using the cytosolic marker (mTurquoise2) and the EL marker (mEmerald), and their stage positions recorded. After labeling, the entire sample insert was carefully removed, and cells were washed three times with photostimulation buffer. After the final wash, photostimulation buffer was replaced with 2 mM MNI in photostimulation buffer, and the sample was returned to the microscope for imaging and photostimulation (see **Photostimulation**). Prior to photostimulation, samples were incubated on the stage for 10 minutes to 1) allow the temperature within the incubator and sample to equilibrate, 2) allow time to re-find the target spines and adjust the xyz positions, and 3) synchronize the start of sLTP after labeling across samples. Images of the target spine and the EL marker were taken before and after sLTP stimulation to ensure successful sLTP induction. After imaging, samples were incubated at 37 °C and 5% CO_2_ for 10 minutes in the stage-top incubator to synchronize the chase step across samples while preparing solutions. Next, MNI was removed and the sample was labeled with 50 nM JF_549_i-HTL in photostimulation buffer (chase) at 37 °C and 5% CO_2_ for 45 minutes on the stage-top incubator. Finally, samples were washed three times in photostimulation buffer and returned to the microscope for imaging. To facilitate the re-identification of stimulated spines after labeling and wash steps, samples were retained in the microscope insert for the duration of the experiment and stage positions were updated accordingly if the sample position shifted. In addition, alignment markers were added to the sample dish and sample insert in case the orientation of the sample was disrupted. Lastly, all labeling and wash steps were performed at a laboratory bench next to the microscope to minimize perturbations to the sample. For pulse-chase experiments involving treatments with pharmacological agents (i.e., latrunculin A, nocodazole, or jasplakinolide), pre-treatments were performed concurrently with JF_635_i-HTL labeling such that the pre-treatment would end at the same time as JF_635_i-HTL labeling. Pharmacological agents were added to all subsequent buffers.

To examine GluA1-HT turnover in response to cLTP, GluA1-HT was labeled with JF_635_i-HTL for 60 minutes at 37 °C and 5% CO_2_ (pulse) and washed three times with chemical stimulation buffer before being stimulated with 0.2 mM glycine for 15 minutes (see **Chemical treatments**). After the induction of cLTP, neurons were rinsed twice with wash buffer and then labeled with JF_549_i-HTL in wash buffer for 45 minutes at 37 °C and 5% CO_2_ (chase). Samples were washed an additional three times in wash buffer prior to imaging. To test the saturation of receptors on the plasma membrane by JF_635_i-HTL, chase labeling with JF_549_i-HTL was performed immediately after removing JF_635_i-HTL and washing the neurons. For this experiment, all steps were performed in ice-cold Hibernate A at 4 °C.

### Image analysis

Colocalization analysis was performed using Imaris imaging software as previously described (Wong et al., 2024). Spine area was measured in Fiji (Schindelin et al., 2012). We first applied an automatic intensity threshold using Otsu’s method to the cytosolic marker (e.g., tdTomato) before and after sLTP stimulation and created binary masks for both conditions. We then isolated the target spine at the base of the neck from rest of the dendrite and determined the spine area before and after stimulation based on the binary masks. To measure the distance between the base of a target spine and the base of its nearest neighboring spine, a straight line between the spine bases was drawn and measured in either the Nikon Elements C Imaging Software or Fiji.

Prior to single-particle tracking (SPT), rolling-ball background subtraction (radius, 25 pixels) and bleach correction (histogram matching) were applied to correct for uneven illumination and loss of signal between time points, respectively. An automatic intensity threshold using Shanbhag’s method was applied to the corrected image and used to subtract background from the image. Finally, we applied a Gaussian filter during single-particle tracking to suppress noise and enhance particles. For display, timelapse images and representative images of EL markers were background-subtracted and bleach-corrected, and thresholded as described for analysis. Display ranges were set identically for Pre and Post images within each experiment. Cytosolic-fill images (tdTomato, GFP, mTurquoise2) were displayed with a contrast range that saturates the dendritic shaft, to show spine morphology. Images were rotated and cropped for presentation; no other adjustments were made. To determine the number of ELs in a specified segment, we counted trajectories in each region after single-particle tracking analysis. For trajectories that occupied more than one specified segment, we attributed the trajectory to the segment that it occupied for most of its duration. For pulse-chase experiments, we used mTurquoise2 as a cytosolic marker.

Although the emission spectrum of mTurquoise2 overlaps with mEmerald, we minimized bleed-through by using appropriate filter sets (CFP em 480/40; GFP em 525/36; Nikon) and imaging sequentially. Moreover, because mEmerald-tagged EL markers form bright puncta on endosomes and lysosomes, we were able to filter out the background created by mTurquoise2 using the automatic intensity threshold. To quantify changes in Tractin intensity, we first subtracted the pixel intensities of the pre-sLTP image from the post-sLTP image to generate an image with pixels that increased in intensity after stimulation. We then applied an automatic intensity threshold (Otsu’s method) to the resulting image to create a mask. We applied this mask to the pre- and post-sLTP images and measured the mean fluorescence intensity (MFI) for Tractin in both cases. To determine the fluorescence intensity of GluA1-HT-JF_635_i and GluA1-HT-JF_549_i, we first created a mask for the dendritic spine using the cytosolic marker (see **above** describing the measurement of spine size). We applied an automatic intensity threshold using the MaxEntropy method, computed separately for each channel, to identify pixels containing GluA1-HT-JF□□□i or GluA1-HT-JF□□□i signal, and measured the MFI of above-threshold pixels of each channel within the target-spine mask. Turnover was calculated by dividing the MFI of JF_549_i (chase) by the MFI of JF_635_i (pulse).

### Single-particle tracking (SPT) and hidden Markov modeling with Bayesian model selection (HMM-Bayes) analysis

Single-particle tracking (SPT) and hidden Markov modeling with Bayesian model selection (HMM-Bayes) were performed with a custom MATLAB pipeline (Wong et al., 2025). Particle localization (x, y) was obtained through 2D Gaussian fitting and tracking was based on the multiple-target tracing (MTT) algorithm (Sergé et al., 2008). The localization and tracking parameters in SPT experiments are listed in Table 1. The resulting tracks were manually curated and inspected to ensure tracks were accurately reconstructed (Wong et al., 2025).

**Table 1.** Localization and tracking parameters used in SLIMfast (MTT program).

| Parameter | Value |
| --- | --- |
| Localization error (for SLIMfast) | 1E-6 |
| Deflation loops | 1 |
| Maximum number of competitors | 3 |
| Maximum diffusion coefficient ($\mu\text{m}^2/\text{s}$) | 0.5 |

The diffusion and transport states of individual endosome and lysosome trajectories were analyzed by the HMM-Bayes MATLAB program using default parameters (Monnier et al., 2015) as previously described (Wong et al., 2024).

### Statistical analysis and experimental replicates

Mixed-effects models, permutation tests, and bootstrap analyses (**Figures 3C**, and **Figures S2D, S11B,** and **S16B**) were performed in Python (v3.11.15) using NumPy (v2.4.4), SciPy (v1.17.1), pandas (v3.0.2), statsmodels (v0.15.0), and patsy (v1.0.3). All other statistical tests were performed in GraphPad Prism (v11.0.2). Specific statistical tests to determine the significance of results are indicated in the figure legends. Statistical analyses were independently verified from the raw data with the assistance of Claude (Anthropic; accessed September 2026), an AI assistant, which also wrote and ran the Python code for analyses in **Figures 3C**, and **Figures S2D, S11B,** and **S16B**. All analyses and results were reviewed by the authors, who take full responsibility for them. Statistical tests involving multiple comparisons (**Figures 1D, 1E**, **2C, 2D**, **3C**, and **3E**; **Figures S2D**, **S3C**, **S4B**, **S5B**, **S6A**, **S6B**, **S6E**, **S7B**, **S10B**, **S12C**, **S13B**, **S13C**, **S13D**, **S14**, **S15B**, **S15C**, and **S16B**) were corrected using the Holm-Šídák step-down method. Within each family of m related comparisons, raw p-values (p_raw_) were ranked in ascending order (rank k = 1 for the smallest p-value). Adjusted p-values (p_adjusted_) were calculated with the following equation:

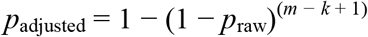

Monotonicity across ranks was enforced by taking the running maximum of adjusted p-values in ascending rank order, ensuring that no lower-ranked comparison received a smaller adjusted p-value than a higher-ranked one. A comparison was considered significant if its adjusted p-value was less than ⍰ = 0.05. Raw p-values are reported in the figure legends; asterisks in the figures reflect the corresponding Holm-Šídák-adjusted p-value. Families were defined as the smallest logically coherent set of comparisons addressing a single question (e.g., one family per experimental condition, per timepoint, or per interaction test, within a given panel).

Colocalization between mEmerald-tagged and endogenous EL markers (**Figure S1**) was evaluated with a non-inferiority test. Colocalization between two endogenous EL markers of the same compartment served as the reference. For each marker, we calculated the difference in mean colocalization coefficients ([GFP AB vs EL marker 2] minus [EL marker 1 vs EL marker 2]) and its one-sided 95% lower confidence bound. The test was one-sided because only reduced colocalization would indicate mislocalization of the mEmerald-tagged marker. Non-inferiority was concluded if the lower bound was greater than -ε, where ε is the non-inferiority margin. ε was set to 0.10 for Pearson coefficients, which is smaller than the range of Pearson coefficients among the four endogenous marker pairs (0.127), and to 0.15 for Manders coefficients. The effect of sLTP on the segment distribution of EL motion (**Figure S2D**) was analyzed with a linear mixed-effects model with timepoint, segment, and their interaction as fixed effects and dendrite as a random intercept, accounting for the fact that all segments and both timepoints derive from the same dendrite. The timepoint × segment interaction was tested with a likelihood-ratio test comparing models with and without the interaction term, both fit by maximum likelihood; model estimates and segment-wise contrasts were obtained from the full model fit by restricted maximum likelihood (REML). Because the fraction of ELs exhibiting active transport is bounded between 0 and 1 and strongly one-inflated before stimulation, each interaction was additionally evaluated with a permutation test in which timepoint labels were shuffled within dendrite (2,000 permutations). Segment-wise differences are reported as the mean change after sLTP with bootstrap 95% confidence intervals (10,000 resamples over dendrites). Interaction tests (**Figure 2**, and **Figures S3C** and **S10B**) were performed to compare the changes (Δ) in paired quantities before and after stimulation between two conditions. In **Figure 2**, we compare changes in the number of ELs (ΔEL#) or fraction of ELs exhibiting active transport (ΔEL AT) before and after sLTP (Post – Pre) between samples that are treated or untreated with latrunculin A (+LatA vs -LatA). In **Figure S3C**, we compare changes in the fraction of ELs exhibiting active transport (ΔEL AT) before and after sLTP stimulation or mock sLTP stimulation in the absence of MNI (+MNI vs -MNI). In **Figure S10B**, we compare the changes in spine area (Δ spine area) immediately after sLTP and 30 minutes after sLTP (sLTP +30 min – sLTP) at spines with ELs present vs ELs absent from their bases (EL^+^ vs EL^-^). A Mann-Whitney test with Holm–Šídák correction was used to determine significance for interaction tests. For experiments where EL motion was imaged after chemical treatments (e.g., cLTP induction), particles were imaged in one segment of a dendrite and then the neuronal culture was discarded. Each time-lapse was derived from an independent neuronal culture to synchronize the start of a time-lapse after treatment across samples. For MNI uncaging experiments, one spine was targeted per neuron (i.e., we identified a spine on a new neuron for each sLTP stimulation experiment). A maximum of 3 spines (i.e., 3 neurons) was targeted for MNI uncaging in one culture before the culture was discarded. Targeted spines were separated by at least 3 mm to minimize the possibility that a spine was exposed to glutamate uncaged in a previous sLTP experiment. For pulse-chase experiments involving sLTP, a maximum of 2 spines was identified per culture prior to stimulation. The two spines were then stimulated in succession to minimize lag between stimulations (see **Pulse-chase labeling with Janelia Fluor dyes**). Because some cultures contributed more than one spine to the +MNI and -MNI turnover comparisons in **Figure 3C**, we additionally fit a linear mixed-effects model (REML; turnover as a function of EL status and EL marker, with a random intercept for culture) to confirm the EL^+^ vs EL^-^ difference was not attributable to spines sharing a common culture. This analysis was not applicable to the +LatA, +Noc, and +Jasp comparisons in **Figure 3**, where each culture contributed only a single spine. This model was also applied to the control vs cLTP comparisons in **Figure S16B**. To test whether EL positioning amplified the sLTP-induced increase in GluA1-HT turnover beyond differences in basal turnover (**Figure S11B**), we extended this model to the combined +MNI and -MNI data from **Figure 3C**, with log-transformed turnover as the response and stimulation (+MNI vs - MNI), EL status, and their interaction as fixed effects (EL marker as a fixed covariate; random intercept for culture). The exponentiated interaction coefficient is the ratio of the sLTP-induced fold-increase in turnover (+MNI relative to -MNI) in EL spines to that in EL^-^ spines, and was tested with a Wald test. Typically, 8–12 transfections were performed after a single dissection (hippocampi pooled from 2–4 rat pups; Wong et al., 2025). 3–4 neuronal cultures were prepared from a single transfection. Experimental replicates for all experiments were derived from at least 3 dissections (≥ 3 biological replicates). We previously demonstrated that variation in vesicle transport behaviors between replicates is less than variation due to treatments (Wong et al., 2024).

## Supporting information

Source Data Main

## Data Availability

All data generated or analyzed during this study are included in the manuscript and supporting files; source data files have been provided for all figures.

## Code Availability

Code and data for the analyses in **Figures 3C, S2D**, **S11B**, and **S16B** are available on Zenodo (https://doi.org/10.5281/zenodo.23073463).

## Acknowledgements

We thank Dr. Luke D. Lavis and Jonathan B. Grimm for providing the Janelia Fluor dyes; Dr. Hyun Ah Yi and the Janelia Research Campus (JRC) Viral Tools facility for packaging AAVs; Dr. Harikrushnan Balasubramanian, Dr. Michael DeSantis, Dr. Jenny Ngo, and the JRC Advance Imaging Center (AIC) for assistance with imaging platforms; Brian Rebhorn and the JRC Media facility for help producing buffers; Phuong Nguyen for help culturing neurons; members of the O’Shea lab for helpful discussions; and Michelle Quiambao for administrative assistance. Claude (Anthropic) was used to assist with proofreading the manuscript text; the authors have reviewed and take responsibility for all content. This work was supported by the Howard Hughes Medical Institute.

## References

Anggono, V., & Huganir, R. L. (2012). Regulation of AMPA receptor trafficking and synaptic plasticity. Current Opinion in Neurobiology, 22(3), 461–469.

Bastin, G., & Heximer, S. P. (2013). Rab family proteins regulate the endosomal trafficking and function of RGS4. The Journal of Biological Chemistry, 288(30), 21836–21849.

Benedetti, L., Barentine, A. E. S., Messa, M., Wheeler, H., Bewersdorf, J., & De Camilli, P. (2018). Light-activated protein interaction with high spatial subcellular confinement. Proceedings of the National Academy of Sciences, 115(10), E2238–E2246.

Benedetti, L., Marvin, J. S., Falahati, H., Guillén-Samander, A., Looger, L. L., & De Camilli, P. (2020). Optimized Vivid-derived Magnets photodimerizers for subcellular optogenetics in mammalian cells. eLife, 9, e63230.

Berry, K. P., & Nedivi, E. (2017). Spine dynamics: Are they all the same? Neuron, 96(1), 43–55.

Bosch, M., Castro, J., Saneyoshi, T., Matsuno, H., Sur, M., & Hayashi, Y. (2014). Structural and molecular remodeling of dendritic spine substructures during long-term potentiation. Neuron, 82(2), 444–459.

Bowen, A. B., Bourke, A. M., Hiester, B. G., Hanus, C., & Kennedy, M. J. (2017). Golgi-independent secretory trafficking through recycling endosomes in neuronal dendrites and spines. eLife, 6, e27362.

Brown, T. C., Correia, S. S., Petrok, C. N., & Esteban, J. A. (2007). Functional compartmentalization of endosomal trafficking for the synaptic delivery of AMPA receptors during long-term potentiation. The Journal of Neuroscience, 27(48), 13311–13315.

Bucci, C., Thomsen, P., Nicoziani, P., McCarthy, J., & van Deurs, B. (2000). Rab7: a key to lysosome biogenesis. Molecular Biology of the Cell, 11(2), 467–480.

Camblor-Perujo, S., & Kononenko, N. L. (2022). Brain-specific functions of the endocytic machinery. The FEBS Journal, 289(8), 2219–2246.

Carroll, R. C., Beattie, E. C., von Zastrow, M., & Malenka, R. C. (2001). Role of AMPA receptor endocytosis in synaptic plasticity. Nature Reviews Neuroscience, 2(5), 315–324.

Chater, T. E., & Goda, Y. (2014). The role of AMPA receptors in postsynaptic mechanisms of synaptic plasticity. Frontiers in Cellular Neuroscience, 8, 401.

Chen, J. W., Murphy, T. L., Willingham, M. C., Pastan, I., & August, J. T. (1985). Identification of two lysosomal membrane glycoproteins. The Journal of Cell Biology, 101(1), 85–95.

Citri, A., & Malenka, R. C. (2008). Synaptic plasticity: Multiple forms, functions, and mechanisms. Neuropsychopharmacology, 33(1), 18–41.

Correia, S. S., Bassani, S., Brown, T. C., Lisé, M. F., Backos, D. S., El-Husseini, A., Passafaro, M., & Esteban, J. A. (2008). Motor protein-dependent transport of AMPA receptors into spines during long-term potentiation. Nature Neuroscience, 11(4), 457–466.

Diering, G. H., & Huganir, R. L. (2018). The AMPA receptor code of synaptic plasticity. Neuron, 100(2), 314–329.

D’Souza, R. S., Semus, R., Billings, E. A., Meyer, C. B., Conger, K., & Casanova, J. E. (2014). Rab4 orchestrates a small GTPase cascade for recruitment of adaptor proteins to early endosomes. Current Biology, 24(11), 1187–1198.

Ehlers, M. D. (2000). Reinsertion or degradation of AMPA receptors determined by activity-dependent endocytic sorting. Neuron, 28(2), 511–525.

Ellis-Davies, G. C. (2007). Caged compounds: photorelease technology for control of cellular chemistry and physiology. Nature Methods, 4(8), 619–628.

Eskelinen, E. L. (2006). Roles of LAMP-1 and LAMP-2 in lysosome biogenesis and autophagy. Molecular Aspects of Medicine, 27(5-6), 495–502.

Esteves da Silva, M., Adrian, M., Schätzle, P., Lipka, J., Watanabe, T., Cho, S., Futai, K., Wierenga, C. J., Kapitein, L. C., & Hoogenraad, C. C. (2015). Positioning of AMPA receptor-containing endosomes regulates synapse architecture. Cell Reports, 13(5), 933–943.

Ferro, L. S., Can, S., Turner, M. A., ElShenawy, M. M., & Yildiz, A. (2019). Kinesin and dynein use distinct mechanisms to bypass obstacles. eLife, 8, e48629.

Fukazawa, Y., Saitoh, Y., Ozawa, F., Ohta, Y., Mizuno, K., & Inokuchi, K. (2003). Hippocampal LTP is accompanied by enhanced F-actin content within the dendritic spine that is essential for late LTP maintenance in vivo. Neuron, 38(3), 447–460.

Goo, M. S., Sancho, L., Slepak, N., Boassa, D., Deerinck, T. J., Ellisman, M. H., Bloodgood, B. L., & Patrick, G. N. (2017). Activity-dependent trafficking of lysosomes in dendrites and dendritic spines. The Journal of Cell Biology, 216(8), 2499–2513.

Grochowska, K. M., Andres-Alonso, M., Karpova, A., & Kreutz, M. R. (2022). The needs of a synapse—how local organelles serve synaptic proteostasis. The EMBO Journal, 41(7), e110057.

Harris, C. R., Millman, K. J., van der Walt, S. J., Gommers, R., Virtanen, P., Cournapeau, D., Wieser, E., Taylor, J., Berg, S., Smith, N. J., Kern, R., Picus, M., Hoyer, S., van Kerkwijk, M. H., Brett, M., Haldane, A., del Río, J. F., Wiebe, M., Peterson, P.,… Oliphant, T. E. (2020). Array programming with NumPy. Nature, 585(7825), 357–362.

Hornfeck, S., Trofimenko, E., & Widmann, C. (2026). Overexpression generates aberrant distribution of endocytic regulators - the case of the Rab11/LAMP1 compartment. PLOS ONE, 21(4), e0346157.

Huganir, R. L., & Nicoll, R. A. (2013). AMPARs and synaptic plasticity: The last 25 years. Neuron, 80(3), 704–717.

Jonker, C. T. H., Deo, C., Zager, P. J., Tkachuk, A. N., Weinstein, A. M., Rodriguez-Boulan, E., Lavis, L. D., & Schreiner, R. (2020). Accurate measurement of fast endocytic recycling kinetics in real time. Journal of Cell Science, 133(2), jcs231225.

Kelly, E. E., Horgan, C. P., McCaffrey, M. W., & Young, P. (2011). The role of endosomal-recycling in long-term potentiation. Cellular and Molecular Life Sciences, 68(2), 185–194.

Kneussel, M., & Hausrat, T. J. (2016). Postsynaptic neurotransmitter receptor reserve pools for synaptic potentiation. Trends in Neurosciences, 39(3), 170–182.

Knödler, A., Feng, S., Zhang, J., Zhang, X., Das, A., Peränen, J., & Guo, W. (2010). Coordination of Rab8 and Rab11 in primary ciliogenesis. Proceedings of the National Academy of Sciences, 107(14), 6346–6351.

Krucker, T., Siggins, G. R., & Halpain, S. (2000). Dynamic actin filaments are required for stable long-term potentiation (LTP) in area CA1 of the hippocampus. Proceedings of the National Academy of Sciences of the United States of America, 97(12), 6856–6861.

Langford, G. M. (2002). Myosin-V, a versatile motor for short-range vesicle transport. Traffic, 3(12), 859–865.

Magee, J. C., & Grienberger, C. (2020). Synaptic plasticity forms and functions. Annual Review of Neuroscience, 43, 95–117.

Man, H. Y., Lin, J. W., Ju, W. H., Ahmadian, G., Liu, L., Becker, L. E., Sheng, M., & Wang, Y. T. (2000). Regulation of AMPA receptor-mediated synaptic transmission by clathrin-dependent receptor internalization. Neuron, 25(3), 649–662.

Matsuzaki, M., Ellis-Davies, G. C., Nemoto, T., Miyashita, Y., Iino, M., & Kasai, H. (2001). Dendritic spine geometry is critical for AMPA receptor expression in hippocampal CA1 pyramidal neurons. Nature Neuroscience, 4(11), 1086–1092.

Matsuzaki, M., Honkura, N., Ellis-Davies, G. C., & Kasai, H. (2004). Structural basis of long-term potentiation in single dendritic spines. Nature, 429(6993), 761–766.

McKinney, W. (2010). Data structures for statistical computing in Python. In S. van der Walt & J. Millman (Eds.), Proceedings of the 9th Python in Science Conference (pp. 56–61).

Monnier, N., Barry, Z., Park, H. Y., Su, K. C., Katz, Z., English, B. P., Dey, A., Pan, K., Cheeseman, I. M., Singer, R. H., & Bathe, M. (2015). Inferring transient particle transport dynamics in live cells. Nature Methods, 12(9), 838–840.

Nakahata, Y., & Yasuda, R. (2018). Plasticity of spine structure: Local signaling, translation and cytoskeletal reorganization. Frontiers in Synaptic Neuroscience, 10, 29.

Nijenhuis, W., van Grinsven, M. M. P., & Kapitein, L. C. (2020). An optimized toolbox for the optogenetic control of intracellular transport. The Journal of Cell Biology, 219(4), e201907149.

Nunez, J. (2008). Primary culture of hippocampal neurons from P0 newborn rats. Journal of Visualized Experiments: JoVE, (19), 895.

Obashi, K., Matsuda, A., Inoue, Y., & Okabe, S. (2019). Precise temporal regulation of molecular diffusion within dendritic spines by actin polymers during structural plasticity. Cell Reports, 27(5), 1503–1515.e8.

Padamsey, Z., McGuinness, L., Bardo, S. J., Reinhart, M., Tong, R., Hedegaard, A., Hart, M. L., & Emptage, N. J. (2017). Activity-dependent exocytosis of lysosomes regulates the structural plasticity of dendritic spines. Neuron, 93(1), 132–146.

Park, M., Penick, E. C., Edwards, J. G., Kauer, J. A., & Ehlers, M. D. (2004). Recycling endosomes supply AMPA receptors for LTP. Science, 305(5692), 1972–1975.

Park, M., Salgado, J. M., Ostroff, L., Helton, T. D., Robinson, C. G., Harris, K. M., & Ehlers, M. D. (2006). Plasticity-induced growth of dendritic spines by exocytic trafficking from recycling endosomes. Neuron, 52(5), 817–830.

Schell, M. J., Erneux, C., & Irvine, R. F. (2001). Inositol 1,4,5-trisphosphate 3-kinase A associates with F-actin and dendritic spines via its N terminus. The Journal of Biological Chemistry, 276(40), 37537–37546.

Schindelin, J., Arganda-Carreras, I., Frise, E., Kaynig, V., Longair, M., Pietzsch, T., Preibisch, S., Rueden, C., Saalfeld, S., Schmid, B., Tinevez, J. Y., White, D. J., Hartenstein, V., Eliceiri, K., Tomancak, P., & Cardona, A. (2012). Fiji: An open-source platform for biological-image analysis. Nature Methods, 9(7), 676–682.

Seabold, S., & Perktold, J. (2010). Statsmodels: Econometric and statistical modeling with Python. In S. van der Walt & J. Millman (Eds.), Proceedings of the 9th Python in Science Conference (pp. 92–96).

Sergé, A., Bertaux, N., Rigneault, H., & Marguet, D. (2008). Dynamic multiple-target tracing to probe spatiotemporal cartography of cell membranes. Nature Methods, 5(8), 687–694.

Somsel Rodman, J., & Wandinger-Ness, A. (2000). Rab GTPases coordinate endocytosis. Journal of Cell Science, 113, 183–192.

Sönnichsen, B., De Renzis, S., Nielsen, E., Rietdorf, J., & Zerial, M. (2000). Distinct membrane domains on endosomes in the recycling pathway visualized by multicolor imaging of Rab4, Rab5, and Rab11. The Journal of Cell Biology, 149(4), 901–914.

Suzuki, K., Tsunekawa, Y., Hernandez-Benitez, R., Wu, J., Zhu, J., Kim, E. J., Hatanaka, F., Yamamoto, M., Araoka, T., Li, Z., Kurita, M., Hishida, T., Li, M., Aizawa, E., Guo, S., Chen, S., Goebl, A., Soligalla, R. D., Qu, J., Jiang, T.,… Izpisua Belmonte, J. C. (2016). In vivo genome editing via CRISPR/Cas9 mediated homology-independent targeted integration. Nature, 540(7631), 144–149.

Swiech, L., Heidenreich, M., Banerjee, A., Habib, N., Li, Y., Trombetta, J., Sur, M., & Zhang, F. (2015). In vivo interrogation of gene function in the mammalian brain using CRISPR-Cas9. Nature Biotechnology, 33(1), 102–106.

Tarantino, N., Tinevez, J. Y., Crowell, E. F., Boisson, B., Henriques, R., Mhlanga, M., Agou, F., Israël, A., & Laplantine, E. (2014). TNF and IL-1 exhibit distinct ubiquitin requirements for inducing NEMO-IKK supramolecular structures. The Journal of Cell Biology, 204(2), 231–245.

Ullrich, O., Reinsch, S., Urbé, S., Zerial, M., & Parton, R. G. (1996). Rab11 regulates recycling through the pericentriolar recycling endosome. The Journal of Cell Biology, 135(4), 913–924.

van Bergeijk, P., Adrian, M., Hoogenraad, C. C., & Kapitein, L. C. (2015). Optogenetic control of organelle transport and positioning. Nature, 518(7537), 111–114.

van Bommel, B., Konietzny, A., Kobler, O., Bär, J., & Mikhaylova, M. (2019). F-actin patches associated with glutamatergic synapses control positioning of dendritic lysosomes. The EMBO Journal, 38(15), e101183.

van der Sluijs, P., Hull, M., Webster, P., Mâle, P., Goud, B., & Mellman, I. (1992). The small GTP-binding protein rab4 controls an early sorting event on the endocytic pathway. Cell, 70(5), 729–740.

Virtanen, P., Gommers, R., Oliphant, T. E., Haberland, M., Reddy, T., Cournapeau, D., Burovski, E., Peterson, P., Weckesser, W., Bright, J., van der Walt, S. J., Brett, M., Wilson, J., Millman, K. J., Mayorov, N., Nelson, A. R. J., Jones, E., Kern, R., Larson, E.,… SciPy 1.0 Contributors. (2020). SciPy 1.0: Fundamental algorithms for scientific computing in Python. Nature Methods, 17(3), 261–272.

Vitelli, R., Santillo, M., Lattero, D., Chiariello, M., Bifulco, M., Bruni, C. B., & Bucci, C. (1997). Role of the small GTPase Rab7 in the late endocytic pathway. The Journal of Biological Chemistry, 272(7), 4391–4397.

Wang, Z., Edwards, J. G., Riley, N., Provance, D. W., Karcher, R., Li, X. D., Davison, I. G., Ikebe, M., Mercer, J. A., Kauer, J. A., & Ehlers, M. D. (2008). Myosin Vb mobilizes recycling endosomes and AMPA receptors for postsynaptic plasticity. Cell, 135(3), 535–548.

Wilkinson, R. S., & Lin, M. Y. (2004). Endocytosis and synaptic plasticity: Might the tail wag the dog? Trends in Neurosciences, 27(4), 171–174.

Wong, V. C., Houlihan, P. R., Liu, H., Walpita, D., DeSantis, M. C., Liu, Z., & O’Shea, E. K. (2024). Plasticity-induced actin polymerization in the dendritic shaft regulates intracellular AMPA receptor trafficking. eLife, 13, e80622.

Wong, V. C., Walpita, D., Liu, Z. J., & O’Shea, E. K. (2025). Single-particle tracking of AMPA receptor-containing vesicles. Bio-protocol, 15(11), e5325.

Wong, V. C., Walpita, D., Liu, Z. J., & O’Shea, E. K. (2026). Code and data for statistical analyses in Figures 3C, S2D, S11B, and S16B (Version 1.0) [Computer software]. Zenodo. 10.5281/zenodo.23073463

Xie, L., Torigoe, S. E., Xiao, J., Mai, D. H., Li, L., Davis, F. P., Dong, P., Marie-Nelly, H., Grimm, J., Lavis, L., Darzacq, X., Cattoglio, C., Liu, Z., & Tjian, R. (2017). A dynamic interplay of enhancer elements regulates *Klf4* expression in naïve pluripotency. Genes & Development, 31(17), 1795–1808.

Yap, C. C., & Winckler, B. (2022). Spatial regulation of endosomes in growing dendrites. Developmental Biology, 486, 5–14.

Yap, C. C., Digilio, L., McMahon, L. P., Garcia, A. D. R., & Winckler, B. (2018). Degradation of dendritic cargos requires Rab7-dependent transport to somatic lysosomes. The Journal of Cell Biology, 217(9), 3141–3159.

