## Supplemental Figures for "Plasticity-induced actin polymerization positions endocytic compartments at the bases of stimulated spines"

Figure S1

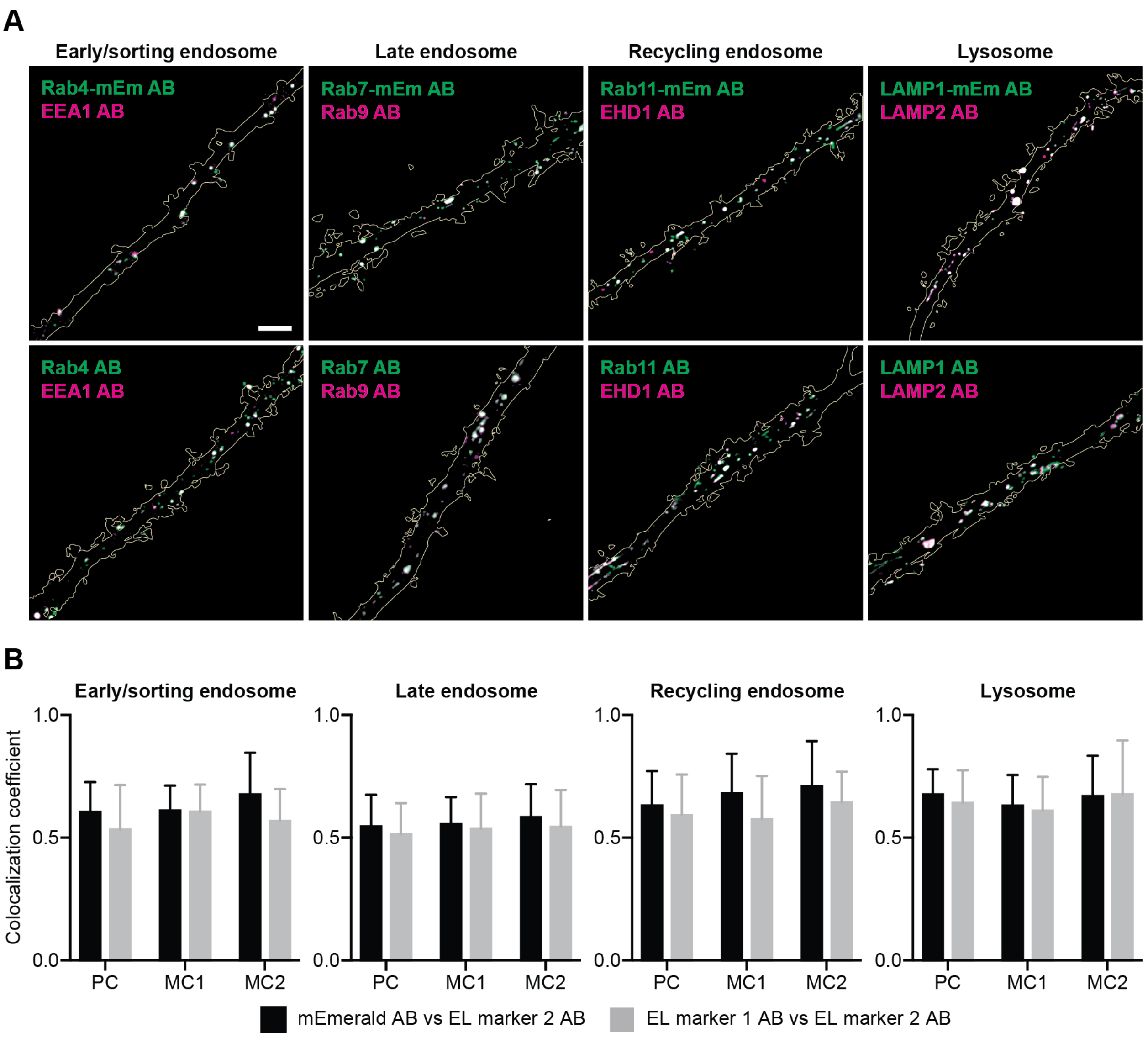
A
Early/sorting endosome
Late endosome
Recycling endosome
Lysosome
Rab4-mEm AB
EEA1 AB
Rab7-mEm AB
Rab9 AB
Rab11-mEm AB
EHD1 AB
LAMP1-mEm AB
LAMP2 AB
Rab4 AB
EEA1 AB
Rab7 AB
Rab9 AB
Rab11 AB
EHD1 AB
LAMP1 AB
LAMP2 AB
B
Early/sorting endosome
Late endosome
Recycling endosome
Lysosome
Colocalization coefficient
1.0
0.5
0.0
PC
MC1
MC2
mEmerald AB vs EL marker 2 AB
EL marker 1 AB vs EL marker 2 AB

**Figure S1. Tagged endosome and lysosome markers expressed from plasmids colocalize with endogenous endosome and lysosome markers.**

(A) Top row: Representative two-color spinning disk confocal images of mEmerald-tagged endosome or lysosome (EL) markers (i.e., Rab4, Rab7, Rab11, and LAMP1) labeled with anti-mEmerald antibody (green) versus antibodies targeting alternative, endogenous EL markers (EEA1, Rab9, EHD1, or LAMP2; magenta). Bottom row: representative two-color spinning disk confocal images of antibodies targeting endogenous Rab4 versus EEA1, Rab7 versus Rab9, Rab11 versus EHD1, and LAMP1 versus LAMP2 in neurons not expressing mEmerald-tagged EL markers (i.e., untransfected neurons). Scale bars, 5 μm.

(B) Colocalization between antibodies targeting mEmerald and EL markers (black) versus antibodies targeting two different EL markers (in neurons not expressing mEmerald-tagged EL markers; gray). Bars represent the mean Pearson correlation coefficients (PC) and Manders overlap coefficients (MC1 and MC2) between antibodies. Error bars represent standard deviation. Colocalization between anti-mEmerald and endogenous marker antibodies was determined to be non-inferior to colocalization between two endogenous marker antibodies for all four EL markers using a non-inferiority margin of 0.10 for the Pearson coefficient (Rab4/EEA1, difference +0.072, one-sided 95% lower bound −0.024; Rab7/Rab9, +0.033, −0.046; Rab11/EHD1, +0.039, −0.056; LAMP1/LAMP2, +0.035, −0.039). The margin was set below the range of Pearson coefficients observed among the four endogenous marker pairs themselves (0.514–0.641, range 0.127). Manders coefficients gave the same result at a margin of

0.15 (see **Methods**). N = 15 spinning disk confocal images derived from 3 labeling experiments for each set of antibodies. Each image is of a dendrite from a different neuron.

### Figure S2

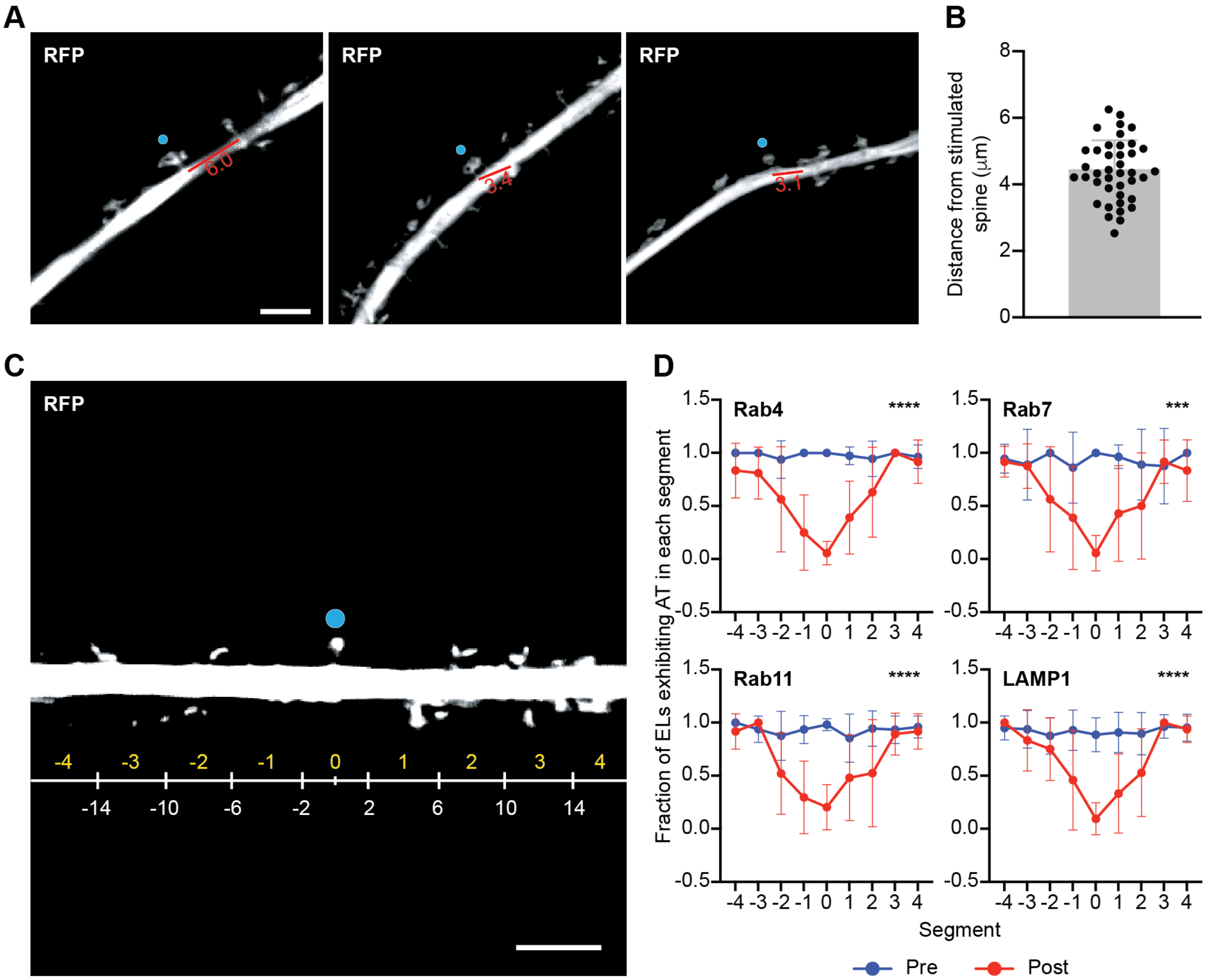
A
RFP
RFP
RFP
6.0
3.4
3.1
B
Distance from stimulated spine (μm)
0
2
4
6
8
C
RFP
-4 -3 -2 -1 0 1 2 3 4
-14 -10 -6 -2 2 6 10 14
D
Rab4 ****
Rab7 ***
Rab11 ****
LAMP1 ****
Fraction of ELs exhibiting AT in each segment
1.5
1.0
0.5
0.0
-0.5
-4 -3 -2 -1 0 1 2 3 4
Segment
Pre
Post

**Figure S2. Distances between sLTP-stimulated spines and neighboring spines.**

(A) Representative spinning disk confocal images of spines selected for structural long-term potentiation (sLTP) stimulation and distances to their nearest neighboring spines. tdTomato (RFP) was used as a cytosolic marker to identify spines and dendritic shafts. Blue dots indicate MNI uncaging sites. Red bars and numbers represent the distance in μm between the bases of targeted spines and the bases of their nearest neighboring spines. Scale bar, 5 μm.

(B) Mean distance between the bases of stimulated spines (i.e., the junction point where a spine neck merges with the dendritic shaft) and the bases of their nearest neighboring spines. N = 40 sLTP stimulation experiments. A new neuron was selected for each sLTP stimulation experiment; a maximum of three spines (i.e., neurons) was stimulated per neuronal culture. We used the mean distance between spine bases, ~4.4 μm (range: 2.5–6.2 μm), to help quantitatively define the region within the dendritic shaft that we consider "near the base" of the stimulated spine. We define "near" as the 4 μm region surrounding the stimulated spine because 2 μm is less than half the distance to the nearest neighboring spine on average.

(C) Representative spinning disk confocal image of a spine and dendrite targeted for sLTP stimulation. tdTomato (RFP) was used as a cytosolic marker to identify spines and dendritic shafts. Blue dot indicates MNI uncaging sites. Dendrite is separated into nine 4 μm segments, labeled in yellow. Hashmark under yellow zero indicates the base of the stimulated spine. Scale bar, 5 μm.

(D) Fraction of mEmerald-tagged $Rab4^+$, $Rab7^+$, or $Rab11^+$ endosomes, or $LAMP1^+$ lysosomes (ELs) that exhibit active transport (AT) in each segment of dendrite (represented by yellow numbers in C) pre-sLTP (Pre, blue) and post-sLTP (Post, red). Bars and error bars represent mean and standard deviation, respectively. Timepoint × segment interaction: Rab4, $\chi^2(8) = 63.7$, ****$p<0.0001$; Rab7, $\chi^2(8) = 30.1$, ***$p=0.0002$; Rab11, $\chi^2(8) = 43.6$, ****$p<0.0001$; LAMP1, $\chi^2(8) = 40.0$, ****$p<0.0001$. Significance determined by a likelihood-ratio test of the timepoint × segment interaction in a linear mixed-effects model with timepoint and segment as fixed effects and dendrite as a random intercept, with Holm–Šídák correction across four EL markers. Each interaction was confirmed by a within-dendrite permutation test (all Holm–Šídák-adjusted $p \leq 0.0020$; see **Methods**). The decrease after sLTP was greatest in the segment containing the base of the stimulated spine ($\Delta = -0.94$ to $-0.78$ across markers; ****$p<0.0001$ for all markers after Holm–Šídák correction). N = 9 sLTP stimulation experiments for each marker. For each experiment, the motion of ELs in the dendrite was imaged before and after sLTP (see **Methods**). Trajectories were binned into each segment and the fraction of trajectories exhibiting active transport in each segment was determined. A new neuron was selected for each sLTP stimulation experiment; a maximum of three spines (i.e., neurons) was stimulated per neuronal culture.

### Figure S3

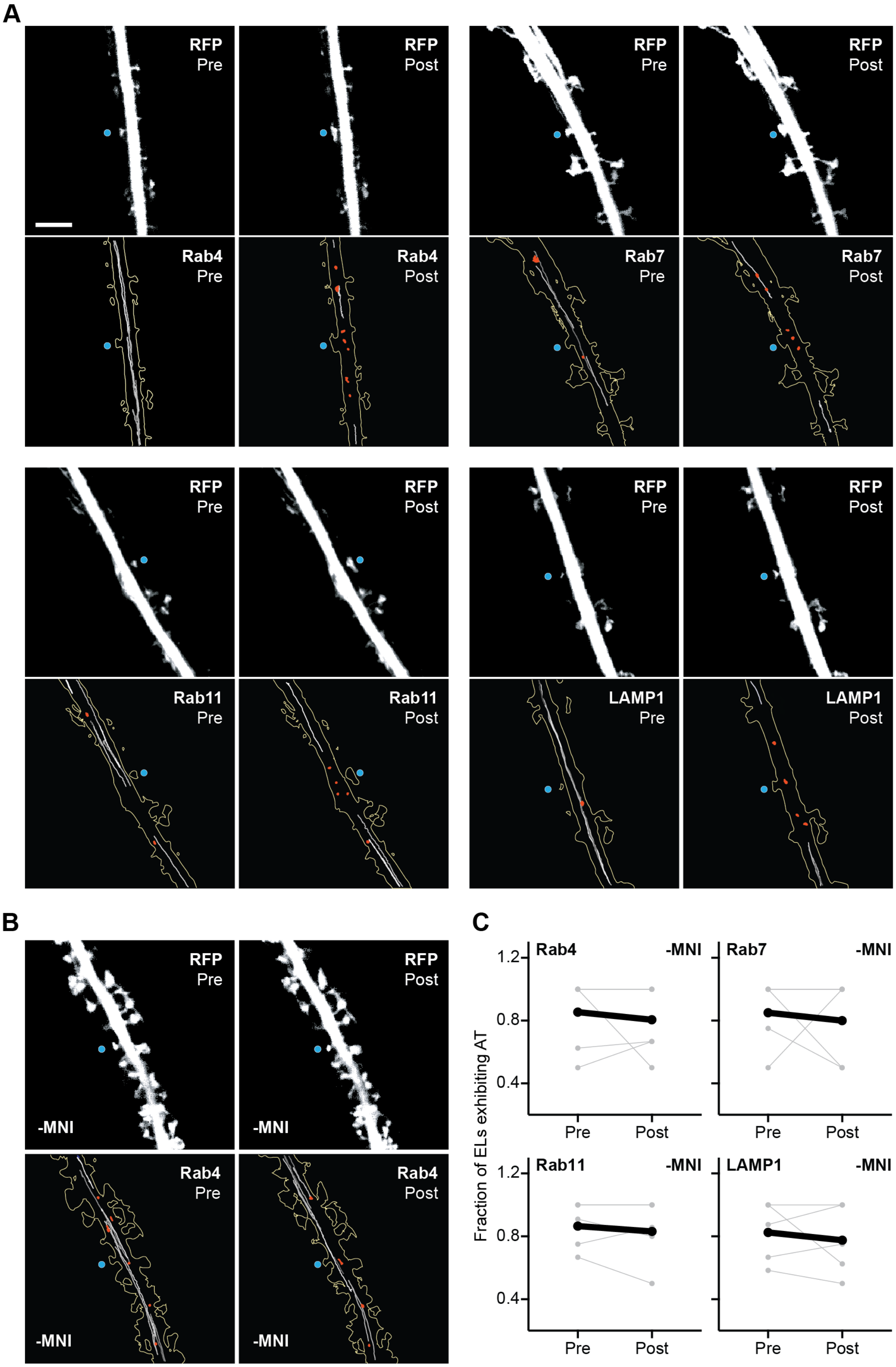
A
RFP
Pre
RFP
Post
Rab4
Pre
Rab4
Post
Rab7
Pre
Rab7
Post
Rab11
Pre
Rab11
Post
LAMP1
Pre
LAMP1
Post
B
-MNI
C
Rab4
Rab7
Rab11
LAMP1
Fraction of ELs exhibiting AT
1.2
0.8
0.4
Pre
Post

**Figure S3. Endosomes and lysosomes lose active transport near the bases of sLTP-stimulated spines.**

(A) RFP (top rows): representative spinning disk confocal or widefield images of dendritic spines expressing tdTomato before and after structural long-term potentiation (sLTP) stimulation (Pre and Post, respectively). Blue dot indicates 4-methoxy-7-nitroindolinyl-caged-L-glutamate (MNI) uncaging site. Scale bar, 5 µm. tdTomato was used to identify and segment the dendritic spine and shaft. Rab#/LAMP1 (bottom rows): trajectories representing the motion of mEmerald-tagged Rab4$^+$, Rab7$^+$, or Rab11$^+$ endosomes, or LAMP1$^+$ lysosomes (ELs) in the dendrites depicted above (RFP, top rows) pre- and post-sLTP. Gray trajectories indicate active transport; red trajectories indicate diffusive motion. Rab4 panels are reproductions of **Figure 1C**.

(B) RFP (top row): representative spinning disk confocal images of a dendritic spine expressing tdTomato before and after mock sLTP stimulation (sLTP stimulation in the absence of MNI, -MNI; Pre and Post, respectively). Rab4 (bottom row): Rab4$^+$ endosome trajectories in the dendrite depicted above (RFP, top row) pre- and post-mock sLTP stimulation.

(C) Fraction of ELs exhibiting active transport (AT) pre- and post-mock sLTP stimulation (-MNI). Each pair of gray dots represents the fraction of ELs exhibiting AT in a dendrite pre- and post-mock sLTP stimulation for one stimulation experiment. Black dots represent the mean fraction of ELs exhibiting AT. Pre-sLTP vs Post-sLTP: Rab4 -MNI, $p>0.9999$; Rab7 -MNI, $p>0.9999$; Rab11 -MNI, $p=0.5000$; LAMP1 -MNI, $p>0.9999$. Significance determined by Wilcoxon signed-rank test with Holm–Šídák correction across four EL markers. Interaction, ΔEL

AT (Post – Pre), +MNI (**Figure 1**) vs -MNI: Rab4, **$p=0.0018$; Rab7, *$p=0.0270$; Rab11, **$p=0.0010$; LAMP1, *$p=0.0070$ (see **Methods**). Significance determined by Mann-Whitney test with Holm–Šídák correction across four EL markers. $N = 5–6$ sLTP stimulation experiments for each marker; for each experiment, the motion of ELs in the dendrite was imaged before and after sLTP (see **Methods**). A new neuron was selected for each sLTP stimulation experiment; a maximum of three spines (i.e., neurons) was stimulated per neuronal culture.

#### Figure S4

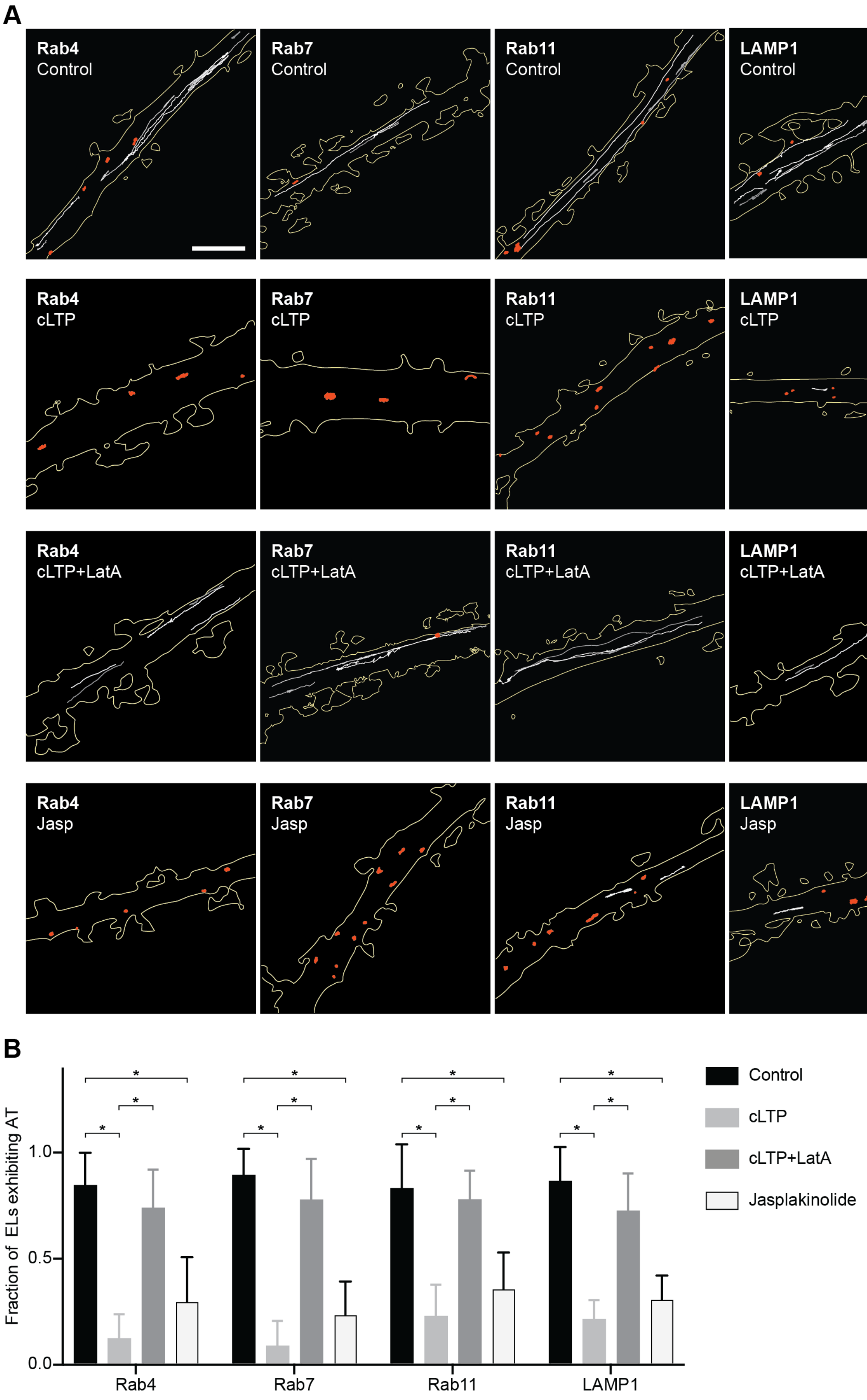


177

**Figure S4. Endosomes and lysosomes lose active transport in response to cLTP and jasplakinolide treatment.**

(A) Trajectories representing the motion of mEmerald-tagged $Rab4^+$, $Rab7^+$, or $Rab11^+$ endosomes, or $LAMP1^+$ lysosomes (ELs) in response to a mock chemical long-term potentiation (cLTP) stimulation (i.e., stimulation buffer without glycine; Control), cLTP stimulation (cLTP), cLTP stimulation in the presence of latrunculin A (cLTP+LatA), and jasplakinolide treatment (Jasp). Gray trajectories indicate active transport; red trajectories indicate diffusive motion. Scale bar, 5 μm. tdTomato was used to identify and segment the dendritic spine and shaft.

(B) Fraction of ELs exhibiting active transport (AT) in response to mock cLTP stimulation (Control), cLTP stimulation (cLTP), cLTP stimulation in the presence of latrunculin A (cLTP+LatA), and jasplakinolide treatment (Jasp). Bars and error bars represent mean and standard deviation, respectively. Rab4: Control vs cLTP, *p=0.0043; cLTP vs cLTP+LatA, *p=0.0079; Control vs Jasp, *p=0.0095. Rab7: Control vs cLTP, *p=0.0043; cLTP vs cLTP+LatA, *p=0.0079; Control vs Jasp, *p=0.0095. Rab11: Control vs cLTP, *p=0.0043; cLTP vs cLTP+LatA, *p=0.0079; Control vs Jasp, *p=0.0095. LAMP1: Control vs cLTP, *p=0.0043; cLTP vs cLTP+LatA, *p=0.0079; Control vs Jasp, *p=0.0095. Significance determined by Mann-Whitney test with Holm–Šídák correction across three comparisons for each marker. N = 4–6 stimulation or treatment experiments for each condition. For each experiment, the motion of ELs was imaged immediately after stimuli or treatments were applied. A new culture of neurons was used for each experiment.

#### Figure S5

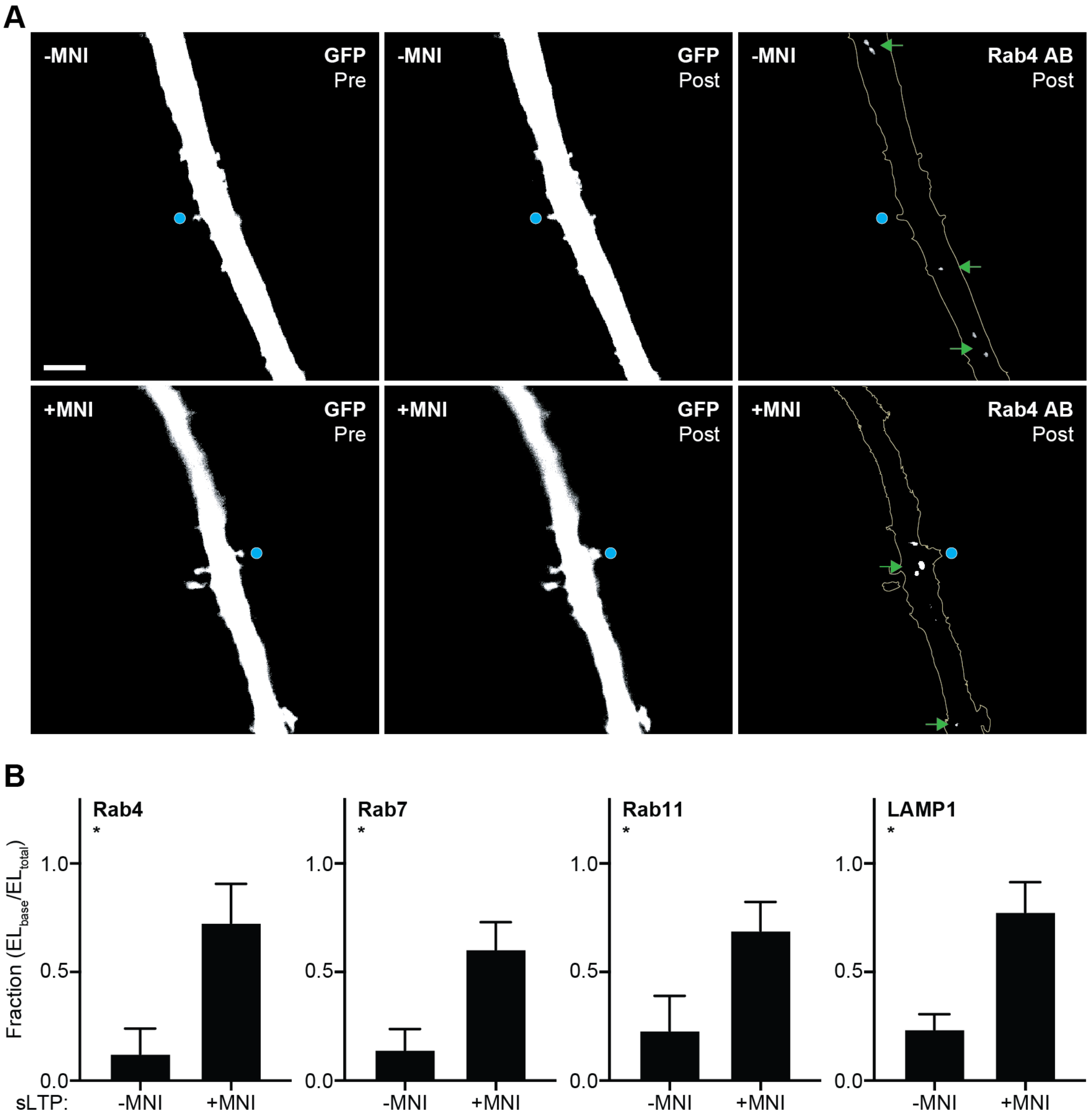


201
202
203
204
205
206

**Figure S5. Endosomes and lysosomes labeled with antibodies are concentrated near the bases of sLTP-stimulated spines.**

(A) GFP (left): representative spinning disk confocal images of dendritic spines expressing GFP before and after structural long-term potentiation (sLTP) stimulation (Pre and Post, respectively) in the absence of MNI (i.e., mock stimulation, -MNI, top row) or the presence of MNI (+MNI, bottom row). Scale bar, 5 μm. Blue dot indicates MNI uncaging site. GFP was used to identify and segment the dendritic spine and shaft. Rab4 AB (right): spinning disk confocal image of the dendrite depicted left (GFP) labeled with an anti-Rab4 antibody after mock sLTP stimulation (-MNI, top row) or sLTP stimulation (+MNI, bottom row). Green arrows indicate endogenous $Rab4^+$ endosomes. Rab4-mEmerald was not expressed in these neurons.

(B) Fraction of endogenous $Rab4^+$, $Rab7^+$, or $Rab11^+$ endosomes, or $LAMP1^+$ lysosomes (ELs) labeled with antibody near the bases of targeted spines (i.e., within the 4 μm region surrounding the stimulated spine base; $EL_{base}/EL_{total}$) after mock sLTP stimulation (-MNI) or sLTP stimulation (+MNI). Bars and error bars represent mean and standard deviation, respectively. Rab4, *p=0.0079; Rab7, *p=0.0286; Rab11, *p=0.0238; LAMP1, *p=0.0079. Significance determined by Mann-Whitney test with Holm–Šídák correction across four EL markers. N = 4–5 sLTP stimulation experiments per marker where images of the spine were taken immediately before and after sLTP, and neurons were fixed immediately after the post-stimulation image. A new neuronal culture was used for each sLTP stimulation experiment. mEmerald-tagged EL markers were not expressed in these neurons.

#### Figure S6

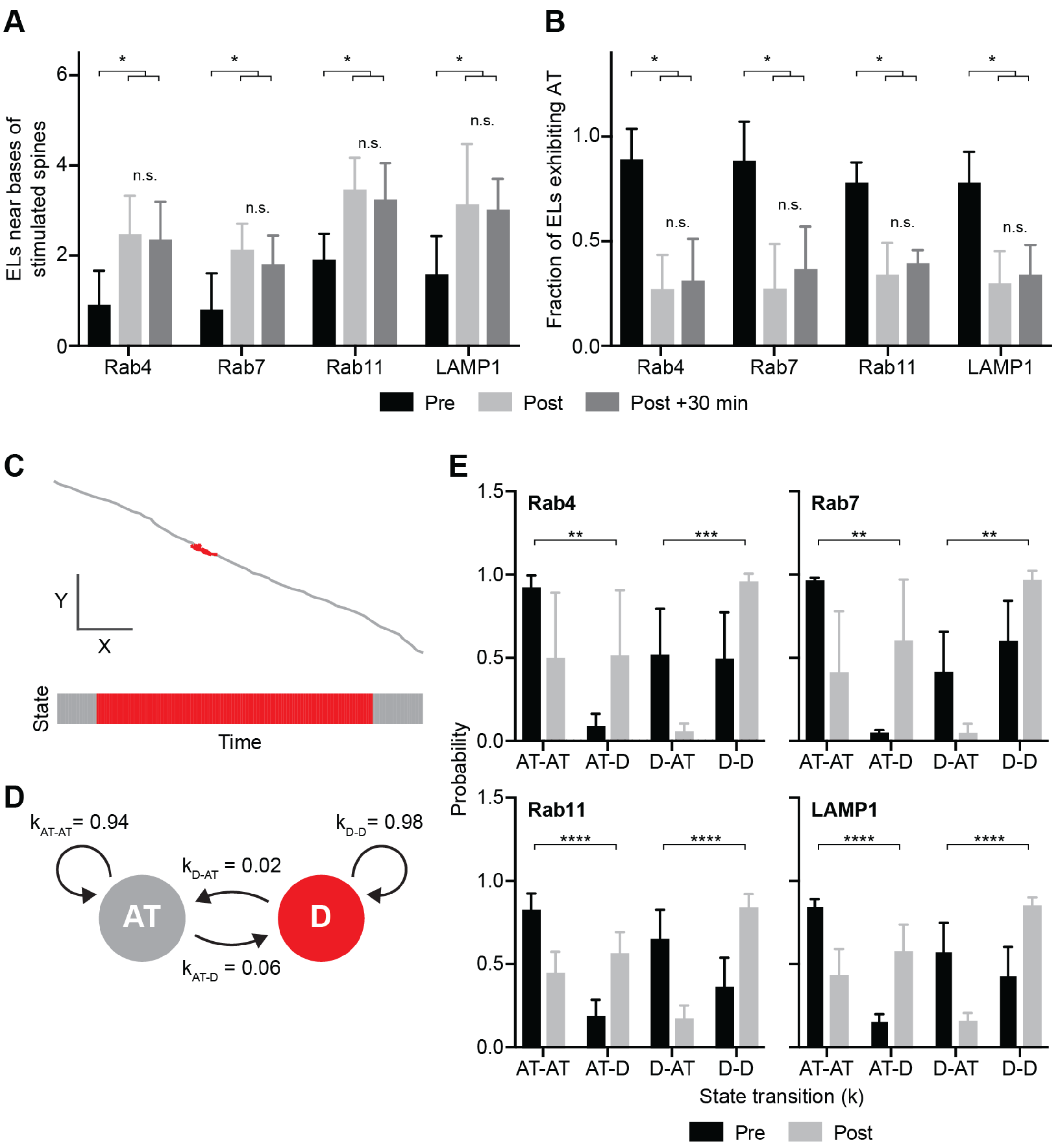

A
B
ELs near bases of stimulated spines
Fraction of ELs exhibiting AT
n.s.
*
Rab4
Rab7
Rab11
LAMP1
Pre
Post
Post +30 min
C
Y
X
State
Time
D
$k_{AT-AT}$= 0.94
$k_{D-D}$= 0.98
$k_{D-AT}$ = 0.02
$k_{AT-D}$ = 0.06
AT
D
E
Probability
**
***
****
AT-AT
AT-D
D-AT
D-D
State transition (k)

**Figure S6. Endosomes and lysosomes are retained near sLTP-stimulated spines.**

(A) Number of mEmerald-tagged Rab4$^+$, Rab7$^+$, or Rab11$^+$ endosomes, or LAMP1$^+$ lysosomes (ELs) near the bases of targeted spines before structural long-term potentiation (sLTP) stimulation (Pre), immediately after sLTP stimulation (Post), or 30 minutes after sLTP stimulation (Post +30 min). Bars and error bars represent mean and standard deviation, respectively. Rab4: Post vs Pre, *p=0.0039; Post +30 min vs Pre, *p=0.0156; Post +30 min vs Post, p>0.9999. Rab7: Post vs Pre, *p=0.0078; Post +30 min vs Pre, *p=0.0156; Post +30 min vs Post, p=0.2500. Rab11: Post vs Pre, *p=0.0078; Post +30 min vs Pre, *p=0.0078; Post +30 min vs Post, p=0.7891. LAMP1: Post vs Pre, *p=0.0078; Post +30 min vs Pre, *p=0.0078; Post +30 min vs Post, p>0.9999. n.s., not significant. Significance determined by Wilcoxon signed-rank test with Holm–Šídák correction across three time-point comparisons for each EL marker. N = 9 sLTP stimulation experiments for each marker. For each experiment, the motion of ELs in the dendrite was imaged immediately before sLTP, immediately after sLTP, and 30 minutes after sLTP and the number of trajectories within a 4 μm segment of dendrite below the stimulated spine was counted for each time point (see **Methods**). A new neuron was selected for each sLTP stimulation experiment; a maximum of three spines (i.e., neurons) was stimulated per neuronal culture.

(B) Fraction of ELs with active transport (AT) before sLTP stimulation (Pre), immediately after sLTP stimulation (Post), or 30 minutes after sLTP stimulation (Post +30 min). Bars and error bars represent mean and standard deviation, respectively. Rab4: Post vs Pre, *p=0.0039; Post +30 min vs Pre, *p=0.0039; Post +30 min vs Post, p=0.4453. Rab7: Post vs Pre, *p=0.0039; Post +30 min vs Pre, *p=0.0078; Post +30 min vs Post, p=0.3281. Rab11: Post vs Pre, *p=0.0039; Post +30 min vs Pre, *p=0.0039; Post +30 min vs Post, p=0.4609. LAMP1: Post vs Pre, *p=0.0039; Post +30 min vs Pre, *p=0.0039; Post +30 min vs Post, p=0.6875. n.s., not significant. Significance determined by Wilcoxon signed-rank test with Holm–Šídák correction across three time-point comparisons for each EL marker. N = 9 sLTP stimulation experiments for each marker. For each experiment, the motion of ELs in the dendrite was imaged immediately before sLTP, immediately after sLTP, and 30 minutes after sLTP. A new neuron was selected for each sLTP stimulation experiment; a maximum of three spines (i.e., neurons) was stimulated per neuronal culture.

(C) Top: exemplar trajectory of an endosome exhibiting both active transport (AT; gray) and diffusion (D; red). Bottom: state sequence of the depicted trajectory (above).

(D) State transition diagram of the depicted trajectory (C) with transition probabilities listed. $k_{AT\text{-}AT}$, remain in active transport; $k_{AT\text{-}D}$, transition from active transport to diffusion; $k_{D\text{-}D}$, remain in diffusion; $k_{D\text{-}AT}$, transition from diffusion to active transport.

(E) State transition probabilities for ELs near the bases of targeted spines pre-sLTP (Pre, black) or post-sLTP (Post, gray). Bars and error bars represent mean and standard deviation, respectively. Rab4, Pre vs Post: $k_{AT\text{-}AT}$ and $k_{AT\text{-}D}$, **p=0.0032; $k_{D\text{-}AT}$ and $k_{D\text{-}D}$, ***p=0.0001. Rab7, Pre vs Post: $k_{AT\text{-}AT}$ and $k_{AT\text{-}D}$, **p=0.0010; $k_{D\text{-}AT}$ and $k_{D\text{-}D}$, **p=0.0010. Rab11, Pre vs Post: $k_{AT\text{-}AT}$ and $k_{AT\text{-}D}$, ****p<0.0001; $k_{D\text{-}AT}$ and $k_{D\text{-}D}$, ****p<0.0001. LAMP1, Pre vs Post: $k_{AT\text{-}AT}$ and $k_{AT\text{-}D}$, ****p<0.0001; $k_{D\text{-}AT}$ and $k_{D\text{-}D}$, ****p<0.0001. Significance determined by Mann-Whitney test with Holm–Šídák correction across two transition comparisons for each marker. For all markers, $k_{D-D} = 1 – k_{D-AT}$ and $k_{AT-AT} = 1 – k_{AT-D}$. N = 5–12 multi-state trajectories pre-sLTP and post-sLTP pooled from 4–6 sLTP stimulation experiments for each EL marker. For each experiment, the motion of ELs in the dendrite was imaged before and after sLTP (see **Methods**). A new neuron was selected for each sLTP stimulation experiment; a maximum of three spines (i.e., neurons) was stimulated per neuronal culture. To avoid issues that might arise from pooling, we also determined significance based on the aggregate k values taken from dendrites (i.e., the mean k values for multi-state trajectories per dendrite). Rab4, Pre vs Post: $k_{AT-AT}$ and $k_{AT-D}$, *p=0.0381; $k_{D-AT}$ and $k_{D-D}$, *p=0.0095. Rab7, Pre vs Post: $k_{AT-AT}$ and $k_{AT-D}$, *p=0.0195; $k_{D-AT}$ and $k_{D-D}$, *p=0.0195. Rab11, Pre vs Post: $k_{AT-AT}$ and $k_{AT-D}$, *p=0.0195; $k_{D-AT}$ and $k_{D-D}$, *p=0.0195. LAMP1, Pre vs Post: $k_{AT-AT}$ and $k_{AT-D}$, *p=0.0079; $k_{D-AT}$ and $k_{D-D}$, *p=0.0079. Significance determined by Mann-Whitney test with Holm–Šídák correction across two transition comparisons for each marker.

#### Figure S7

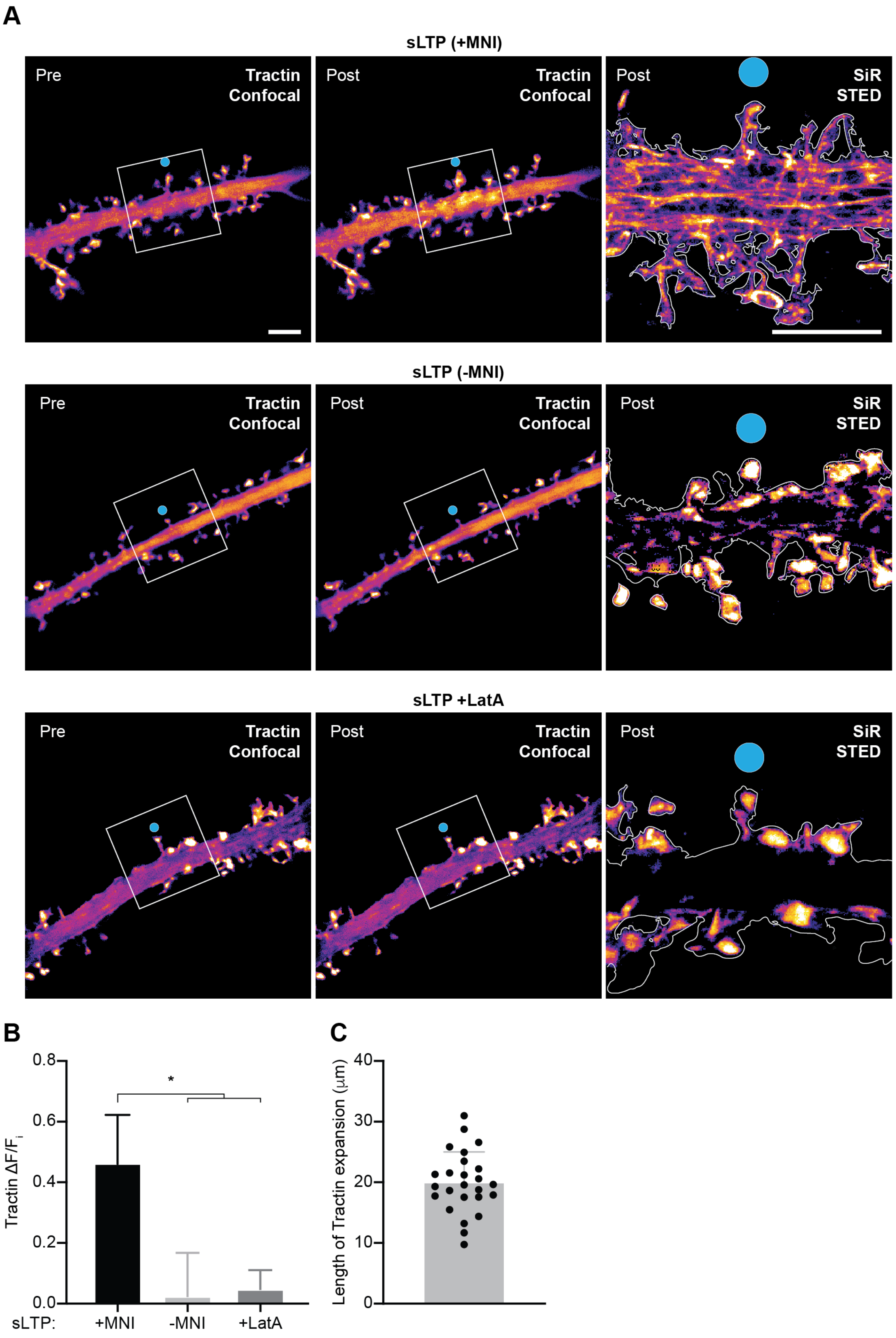


305

**Figure S7. F-actin networks form at the bases of sLTP-stimulated spines.**

(A) Tractin (confocal): representative spinning disk confocal images of F-tractin-mNeonGreen (Tractin) before and after structural long-term potentiation (sLTP) stimulation (Pre and Post, respectively) in the presence of MNI (+MNI, top), the absence of MNI (i.e., mock, -MNI, center), or in the presence of latrunculin A (+LatA, bottom). Scale bar, 5 μm. SiR (STED): stimulated emission depletion (STED) images of the highlighted region (white box) after sLTP (+MNI, top), mock sLTP stimulation in the absence of MNI (-MNI, center), or sLTP stimulation in the presence of latrunculin A (+LatA, bottom) labeled with silicon rhodamine (SiR)-actin. Scale bar, 5 μm. Blue dot indicates MNI uncaging site.

(B) Increase in F-tractin-mScarlet (Tractin) mean fluorescence intensity (MFI; $\Delta F/F_i$) after sLTP (+MNI), mock sLTP stimulation in the absence of MNI (-MNI), or sLTP stimulation in the presence of latrunculin A (+LatA). Bars and error bars represent mean and standard deviation, respectively. +MNI vs -MNI, *p=0.0079; +MNI vs +LatA, *p=0.0079. Significance determined by Mann-Whitney test with Holm–Šídák correction across the comparisons shown in graph. N = 5 sLTP stimulation experiments for each condition where Tractin was imaged before and after sLTP. A new neuron was selected for each sLTP stimulation experiment; a maximum of two spines (i.e., neurons) was stimulated per neuronal culture.

(C) Length of dendrite exhibiting increased F-tractin-mScarlet (Tractin) intensity. Bars and error bars represent mean and standard deviation, respectively. N = 25 sLTP stimulation experiments where Tractin was imaged before and after sLTP. A new neuron was selected for each sLTP stimulation experiment; a maximum of three spines (i.e., neurons) was stimulated per neuronal culture. We used the mean length of Tractin expansion, ~20 μm, to define approximately the region where there would be actin polymerization in response to sLTP if latrunculin A treatment did not prevent actin polymerization.

**Figure S8**

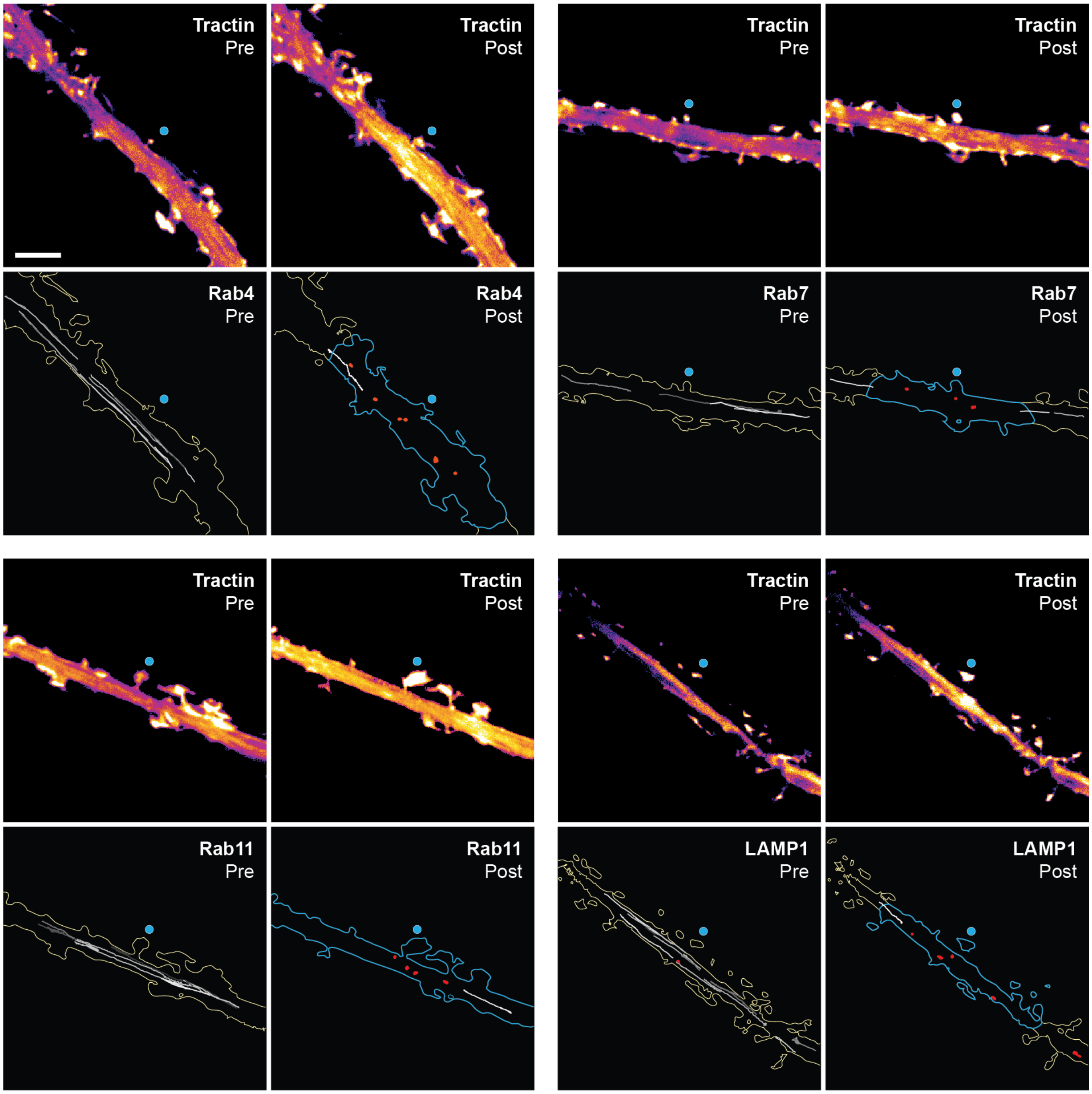


352

353

354

355

356

**Figure S8. Endosomes and lysosomes lose active transport inside regions with increased F-actin.**

Tractin (top rows): representative spinning disk confocal or widefield images of dendrites expressing F-tractin-mScarlet before and after structural long-term potentiation (sLTP) stimulation (Pre and Post, respectively). Blue dot indicates MNI uncaging site. Scale bar, 5 µm. Rab#/LAMP1 (bottom rows): trajectories representing the motion of mEmerald-tagged $Rab4^+$, $Rab7^+$, or $Rab11^+$ endosomes, or $LAMP1^+$ lysosomes (ELs) in the dendrites depicted above (Tractin, top rows) pre- and post-sLTP. Blue outline indicates region with increased Tractin signal after sLTP. Gray trajectories indicate active transport; red trajectories indicate diffusive motion. Rab4 panels are reproductions of **Figure 2A-B**.

#### Figure S9

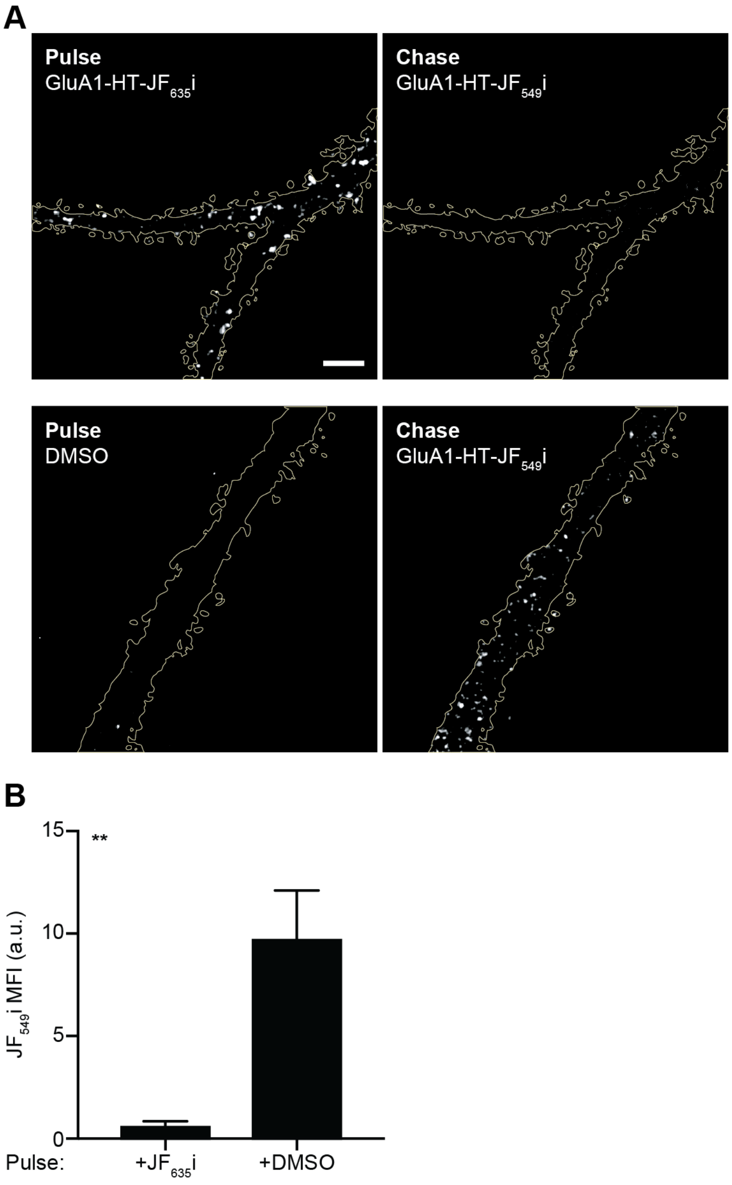
A
Pulse
GluA1-HT-JF635i
Chase
GluA1-HT-JF549i
Pulse
DMSO
Chase
GluA1-HT-JF549i
B
JF549i MFI (a.u.)
15
10
5
0
**
Pulse:
+JF635i
+DMSO

**Figure S9. $JF_{635}$i-HTL labeling blocks most surface GluA1-HT receptors.**

(A) Representative spinning disk confocal images of dendrites expressing GluA1-HT first blocked with $JF_{635}$i-HTL (top left) or DMSO (i.e., no $JF_{635}$i-HTL control; bottom left) and then chased with $JF_{549}$i-HTL (right). Scale bar, 5 μm. GFP was used to identify and segment the dendritic spine and shaft. To test whether $JF_{635}$i-HTL could block the majority of surface receptors, all steps of this experiment were performed at 4 °C using cold buffers to inhibit exocytosis of receptors during and after blocking.

(B) Mean fluorescence intensity (MFI) of GluA1-HT labeled with $JF_{549}$i-HTL (chase) after blocking with $JF_{635}$i-HTL or DMSO. a.u., arbitrary units. Bars and error bars represent mean and standard deviation, respectively. **p=0.0022 by Mann-Whitney test. N = 6 dendrite images derived from three labeling experiments for each condition.

#### Figure S10

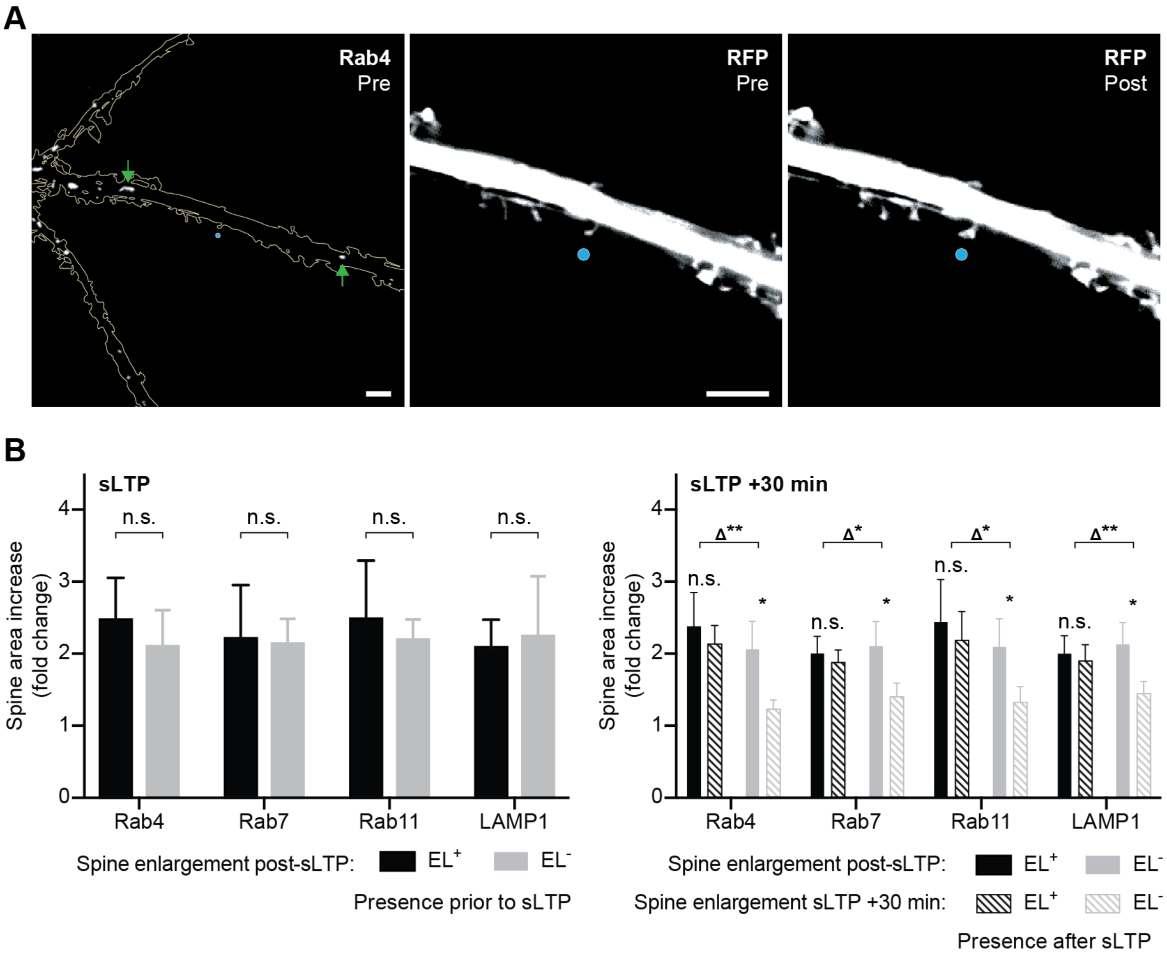
A
Rab4
Pre
RFP
Pre
RFP
Post
B
sLTP
n.s.
Spine area increase
(fold change)
Rab4
Rab7
Rab11
LAMP1
Spine enlargement post-sLTP:
EL+
EL-
Presence prior to sLTP
sLTP +30 min
Δ**
Δ*
n.s.
*
Spine enlargement sLTP +30 min:
Presence after sLTP

**Figure S10. Endosome and lysosome positioning is not required for initial spine enlargement in response to sLTP.**

(A) Rab4 (left): representative spinning disk confocal image of Rab4-mEmerald$^+$ (Rab4$^+$) endosomes in a dendrite prior to structural long-term potentiation (sLTP) stimulation. Green arrows indicate Rab4$^+$ endosomes. RFP (right): dendritic spine expressing tdTomato on the dendrite depicted left (Rab4, left) before and after sLTP (Pre and Post, respectively). Scale bars, 5 μm. Blue dot indicates MNI uncaging site.

(B) Left: increase in spine area (fold change) immediately after sLTP stimulation (sLTP) for spines with mEmerald-tagged Rab4$^+$, Rab7$^+$, or Rab11$^+$ endosomes, or LAMP1$^+$ lysosomes (ELs) present at their bases (EL$^+$, black) or absent from their bases (EL$^-$, gray) prior to stimulation. Bars and error bars represent mean and standard deviation, respectively. Rab4, p=0.3282; Rab7, p=0.9591; Rab11, p=0.7209; LAMP1, p=0.9591. n.s., not significant. Significance determined by Mann-Whitney test with Holm–Šídák correction across four EL markers. N = 16 sLTP stimulation experiments for each EL marker (i.e., 8 sLTP experiments for spines with ELs present and 8 sLTP experiments for spines without ELs present). For each experiment, the target spine was imaged immediately before and after sLTP stimulation. A new neuron was selected for each sLTP stimulation experiment; a maximum of three spines (i.e., neurons) was stimulated per neuronal culture. Right: increase in spine area (fold change) 30 minutes after sLTP stimulation (sLTP +30 min) for spines with ELs present at their bases (EL$^+$, black stripes) or absent from their bases (EL$^-$, gray stripes) after stimulation. Bars and error bars represent mean and standard deviation, respectively. EL$^+$, sLTP vs sLTP +30 min: Rab4, p=0.1094; Rab7, p=0.1094; Rab11, p=0.0781; LAMP1, p=0.1484. n.s., not significant. $EL^-$, sLTP vs sLTP +30 min: Rab4, *p=0.0078; Rab7, *p=0.0078; Rab11, *p=0.0078; LAMP1, *p=0.0078. Significance determined by Wilcoxon signed-rank test with Holm–Šídák correction across four EL markers. Interaction, Δ spine area (sLTP +30 min – sLTP), $EL^+$ vs $EL^-$: Rab4, **p=0.0019; Rab7, *p=0.0148; Rab11, *p=0.0207; LAMP1, **p=0.0003 (See **Methods**). Significance determined by Mann-Whitney test with Holm–Šídák correction across four EL markers. N = 16 sLTP stimulation experiments for each EL marker (i.e., 8 uncaging experiments for spines with ELs present and 8 uncaging experiments for spines without ELs present). For each experiment, the target spine was imaged immediately before sLTP stimulation, immediately after sLTP stimulation, and 30 minutes after sLTP stimulation. A new neuron was selected for each sLTP stimulation experiment; a maximum of two spines (i.e., neurons) was stimulated per neuronal culture. **Figure S10B**, left and right are different sets of experiments.

#### Figure S11

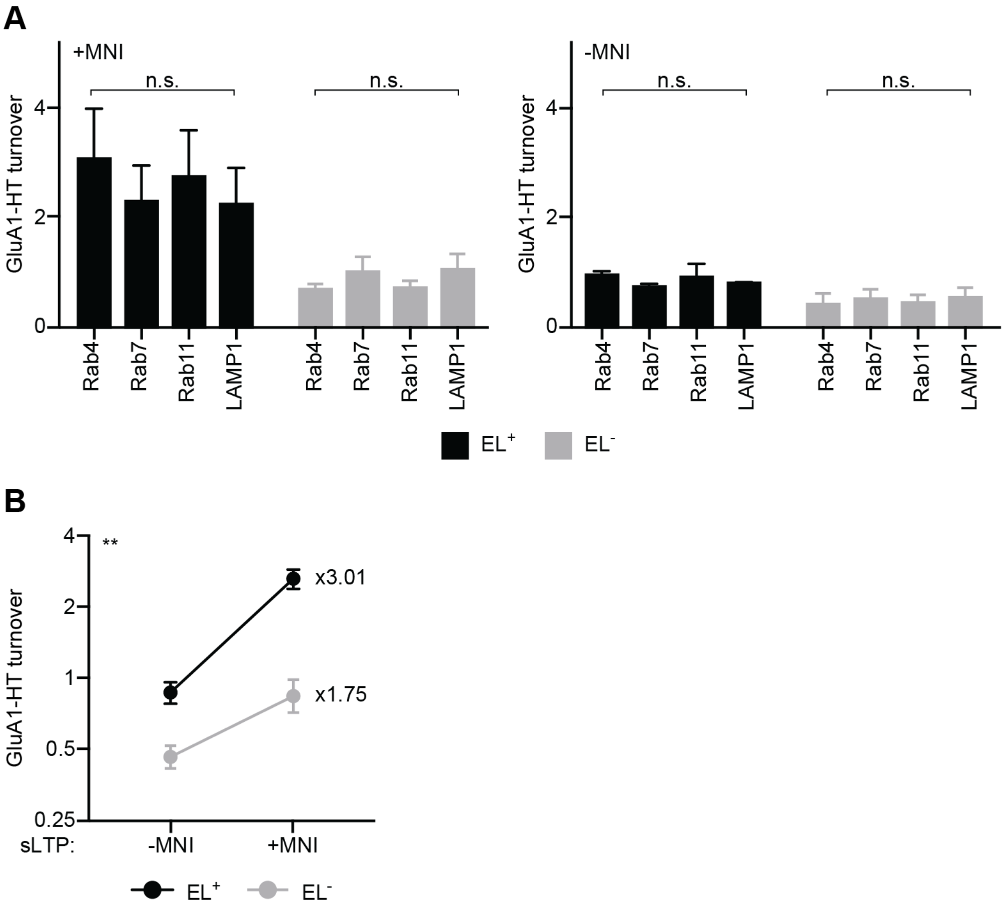

A
+MNI
-MNI
n.s.
GluA1-HT turnover
0
2
4
Rab4
Rab7
Rab11
LAMP1
EL+
EL-
B
**
x3.01
x1.75
4
2
1
0.5
0.25
sLTP:
-MNI
+MNI

**Figure S11. GluA1-HT turnover in response to sLTP depends on endosome or lysosome positioning and is consistent across markers.**

(A) +MNI (left): GluA1-HT turnover in response to structural long-term potentiation (sLTP) stimulation on spines at which there are mEmerald-tagged $Rab4^{+}$, $Rab7^{+}$, or $Rab11^{+}$ endosomes, or $LAMP1^{+}$ lysosomes (ELs) present ($EL^{+}$, black, left) or absent ($EL^{-}$, gray, right) near their bases after sLTP stimulation. Bars and error bars represent mean and standard deviation, respectively. $EL^{+}$ (black), p=0.0697; $EL^{-}$ (gray), p=0.0926. n.s., not significant. N = 14–16 sLTP stimulation experiments for each EL marker. For each experiment, images of the spine and EL marker were taken immediately before and after sLTP stimulation, while images of the EL marker and pulse-chase labeling were taken after the chase (see **Methods**). -MNI (right): GluA1-HT turnover in response to mock sLTP stimulation in the absence of MNI on spines at which there are ELs present ($EL^{+}$, black, left) or absent ($EL^{-}$, gray, right) near their bases after mock stimulation. Bars and error bars represent mean and standard deviation, respectively. $EL^{+}$ (black), p=0.1748; $EL^{-}$ (gray), p=0.2125. N = 14–15 sLTP stimulation experiments for each EL marker. Significance determined by Kruskal-Wallis test. Because turnover shows similar trends in response to stimulation across all four markers, we pooled together the turnover data from each marker to increase statistical power of results for **Figure 3**.

(B) GluA1-HT turnover after mock sLTP stimulation (-MNI) or sLTP stimulation (+MNI) in spines at which ELs are present ($EL^{+}$, black) or absent ($EL^{-}$ gray) near their bases after stimulation. Dots and error bars represent mean and 95% confidence interval, respectively. Lines connect the means for $EL^{+}$ and $EL^{-}$ spines. Numbers indicate the fold-increase in mean turnover after sLTP relative to mock stimulation for spines with the same EL status. The y-axis is plotted on a $\log_2$ scale. Stimulation × EL interaction: ratio of fold-increases ($EL^+/EL^-$) = 1.63 (95% CI 1.19–2.23), **p=0.0024. Significance determined by a linear mixed-effects model on log-transformed turnover with stimulation, EL status, and their interaction as fixed effects, EL marker as a fixed covariate, and culture as a random intercept (see **Methods**). N = 12–47 sLTP- or mock-stimulated spines pooled from Rab4, Rab7, Rab11, and LAMP1 experiments. ELs were absent from the bases of all target spines prior to stimulation. Turnover data are a reproduction of data from **Figure 3C**, bottom. A new neuron was selected for each sLTP stimulation experiment; a maximum of two spines (i.e., neurons) was stimulated per neuronal culture. This analysis accounts for differences in basal turnover between spines with and without EL positioning; a ratio greater than 1 indicates that EL positioning is associated with a larger sLTP-induced fold-increase in GluA1-HT turnover.

#### Figure S12

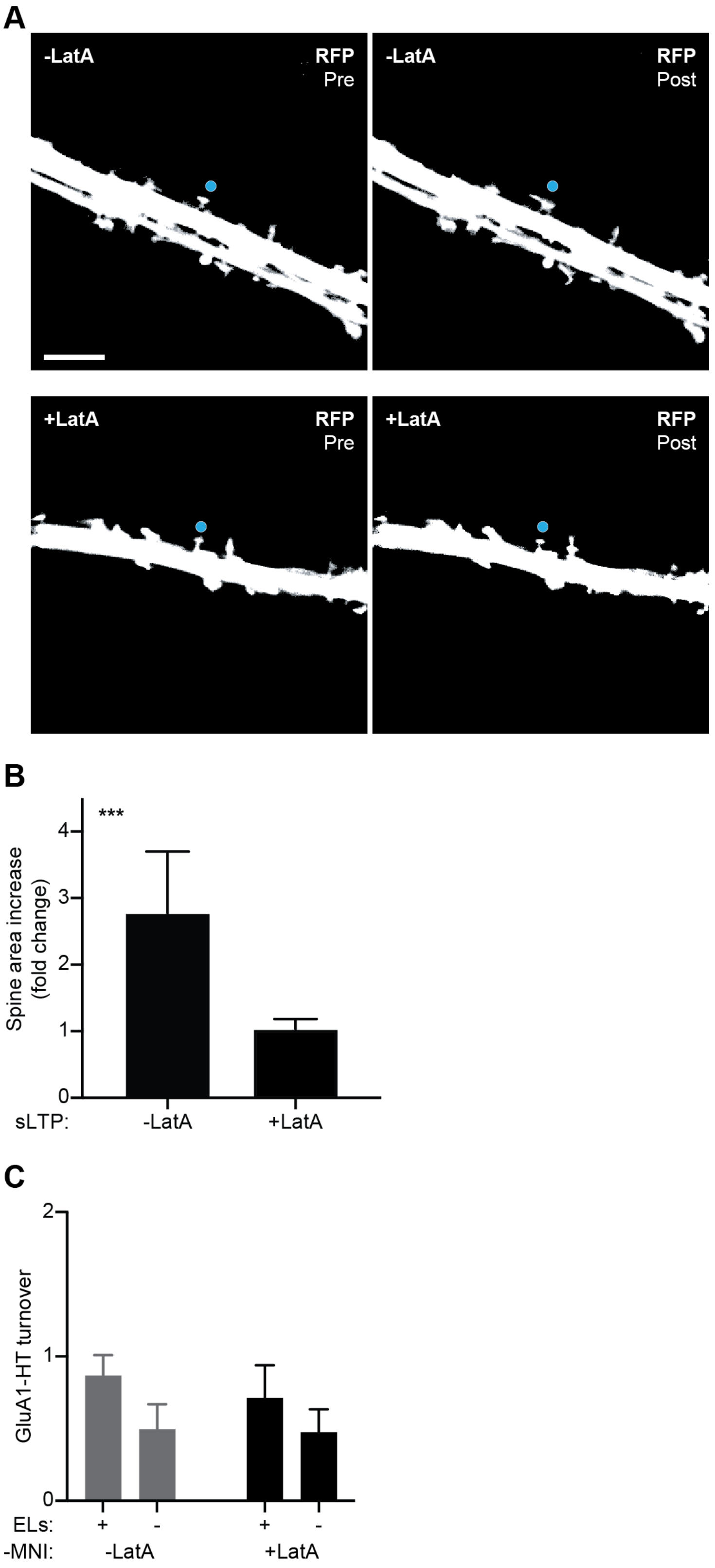
A
-LatA
RFP
Pre
-LatA
RFP
Post
+LatA
RFP
Pre
+LatA
RFP
Post
B
***
Spine area increase (fold change)
0
1
2
3
4
sLTP:
-LatA
+LatA
C
GluA1-HT turnover
0
1
2
ELs:
+
-
+
-
-MNI:
-LatA
+LatA

**Figure S12. Latrunculin A treatment inhibits spine enlargement in response to sLTP but not GluA1-HT turnover in response to mock sLTP stimulation.**

(A) Representative confocal images of dendrites expressing tdTomato pre-sLTP and post-sLTP in the absence of latrunculin A (-LatA, top row) or presence of latrunculin A (+LatA, bottom row). Blue dot indicates MNI uncaging site. Scale bar, 5 μm.

(B) Increase in spine area (fold change) after sLTP stimulation in the absence of latrunculin A (-LatA) or presence of latrunculin A (+LatA). Bars and error bars represent mean and standard deviation, respectively. ***p=0.0002 by Mann-Whitney test. N = 7–9 sLTP stimulation experiments for each condition. For each experiment, the target spine was imaged immediately before and after sLTP stimulation. A new neuron was selected for each sLTP stimulation experiment; a maximum of three spines (i.e., neurons) was stimulated per neuronal culture.

(C) GluA1-HT turnover in response to mock structural long-term potentiation (sLTP) stimulation (-MNI) in the absence or presence of latrunculin A (-LatA, gray or +LatA, black, respectively) on spines at which there are mEmerald-tagged $Rab4^+$, $Rab7^+$, or $Rab11^+$ endosomes, or $LAMP1^+$ lysosomes present or absent (EL: +/-) near their bases after mock stimulation. Bars and error bars represent mean and standard deviation, respectively. -LatA $EL^+$ vs +LatA $EL^+$, p=0.2615; -LatA $EL^-$ vs +LatA $EL^-$, p=0.7402; +LatA $EL^+$ vs +LatA $EL^-$, p=0.0418. Significance determined by Mann-Whitney test with Holm–Šídák correction across the comparisons shown in graph. N = 16–58 mock sLTP-stimulated spines for each condition pooled from 4–12 experiments each for Rab4, Rab7, Rab11, or LAMP1. For pulse-chase experiments analyzed in (C), we applied mock stimulation to dendritic spines that did not have ELs near their bases prior to mock stimulation. For each experiment, images of the spine and EL marker were taken immediately before and after mock sLTP stimulation, while images of the EL marker and pulse-chase labeling were taken after the chase (see **Methods**). -LatA $EL^+$ and $EL^-$ turnover data are a reproduction of the turnover data from **Figure 3C**, -MNI $EL^+$ and $EL^-$.

#### Figure S13

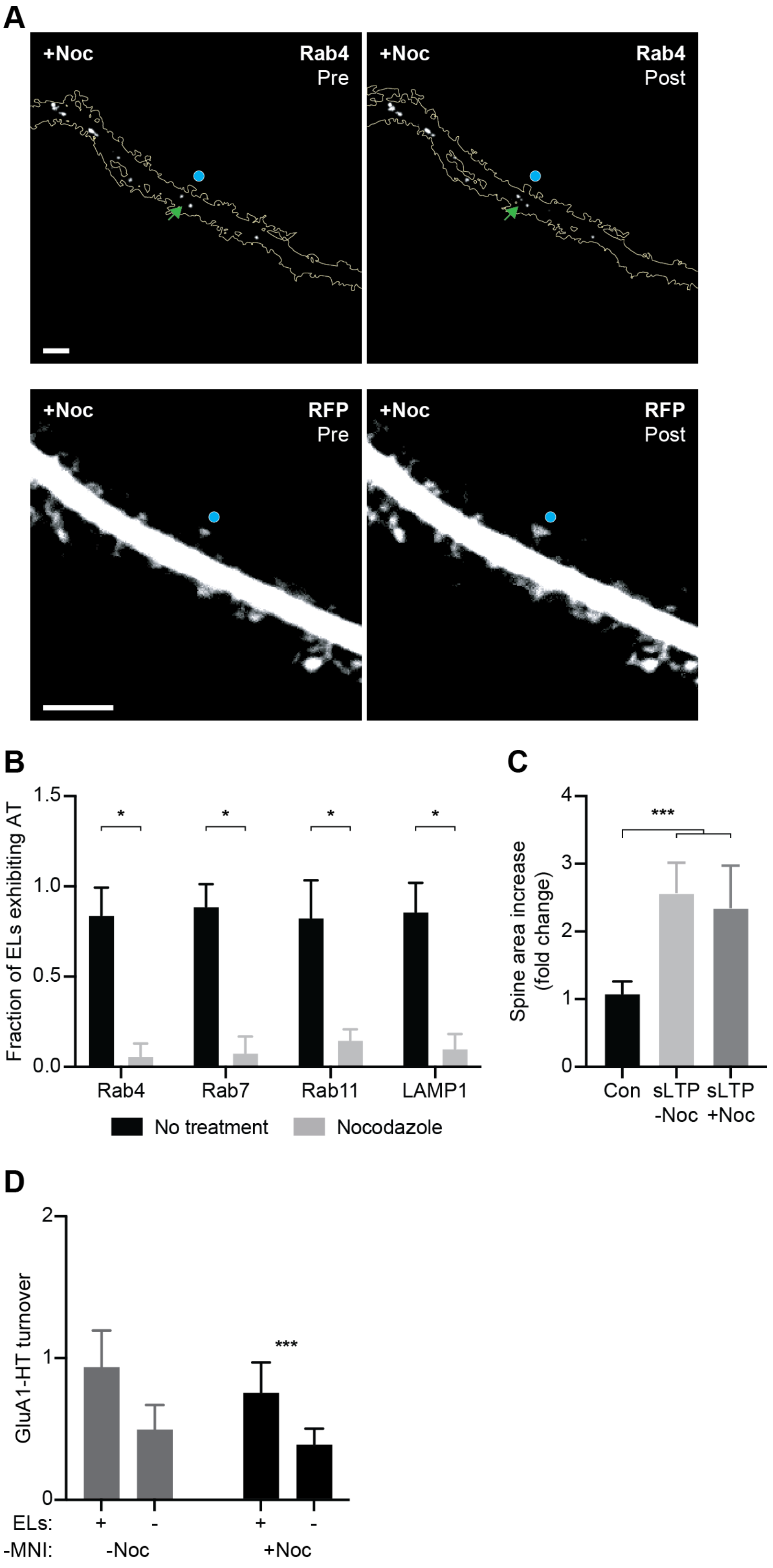


565

**Figure S13. Nocodazole treatment inhibits transport of endosomes and lysosomes but not spine enlargement in response to sLTP or GluA1-HT turnover in response to mock sLTP stimulation.**

(A) Rab4 (top): representative spinning disk confocal images of Rab4-mEmerald$^+$ (Rab4$^+$) endosomes before and after structural long-term potentiation (sLTP) stimulation (Pre and Post, respectively) in the presence of nocodazole (+Noc). Green arrows indicate Rab4$^+$ endosomes. RFP (bottom): dendritic spine depicted above (Rab4, top) expressing tdTomato pre-sLTP (Pre) and post-sLTP (Post) in the presence of nocodazole. Scale bars, 5 μm. Blue dot indicates MNI uncaging site.

(B) Fraction of mEmerald-tagged Rab4$^+$, Rab7$^+$, or Rab11$^+$ endosomes, or LAMP1$^+$ lysosomes (ELs) with active transport after treatment with nocodazole. Bars and error bars represent mean and standard deviation, respectively. Rab4, *p=0.0095; Rab7, *p=0.0095; Rab11, *p=0.0095; LAMP1, *p=0.0095. Significance determined by Mann-Whitney test with Holm–Šídák correction across four EL markers. N = 4–6 treatment experiments for each condition; for each experiment, the motion of ELs in a dendrite was imaged immediately after treatment. A new neuronal culture was used for each experiment. The no treatment control data are a reproduction of the mock chemical long-term potentiation (cLTP) data from **Figure S4**.

(C) Increase in spine area (fold change) after sLTP in the absence of nocodazole (-Noc) or sLTP in the presence of nocodazole (+Noc). Bars and error bars represent mean and standard deviation, respectively. For control (Con), mock sLTP stimulation was performed without MNI (i.e., -MNI). Con vs sLTP -Noc, ***p=0.0002; Con vs sLTP +Noc, ***p=0.0007. Significance determined by Mann-Whitney test with Holm–Šídák correction across the comparisons shown in graph. N = 6–8 sLTP experiments for each condition; for each experiment, the target spine was imaged immediately before and after sLTP stimulation. A new neuron was selected for each sLTP stimulation experiment; a maximum of three spines (i.e., neurons) was stimulated per neuronal culture.

(D) GluA1-HT turnover in response to mock sLTP stimulation (-MNI) in the absence or presence of nocodazole (-Noc, gray, or +Noc, black, respectively) on spines at which there are ELs present or absent (EL: +/-) near their bases before and after mock stimulation. Bars and error bars represent mean and standard deviation, respectively. -Noc $EL^+$ vs +Noc $EL^+$, p=0.1442; -Noc $EL^-$ vs +Noc $EL^-$, p=0.0529; +Noc $EL^+$ vs +Noc $EL^-$, ***p=0.0003. Significance determined by Mann-Whitney test with Holm–Šídák correction across the comparisons shown in graph. N = 19–58 mock sLTP-stimulated spines for each condition pooled from 4–12 experiments each for Rab4, Rab7, Rab11, or LAMP1. For pulse-chase experiments analyzed in (D), we applied mock stimulation to dendritic spines that either did or did not have ELs near their bases prior to mock stimulation. EL positioning did not change in response to mock stimulation. For each experiment, images of the spine and EL marker were taken immediately before and after mock sLTP stimulation, while images of the EL marker and pulse-chase labeling were taken after the chase (see **Methods**). A new neuronal culture was used for each experiment. -Noc $EL^+$ data is distinct from data from **Figure 3C**, -MNI $EL^+$ (ELs are positioned prior to mock stimulation). -Noc $EL^-$ turnover data is a reproduction of turnover data from **Figure 3C**, -MNI $EL^-$.

#### Figure S14

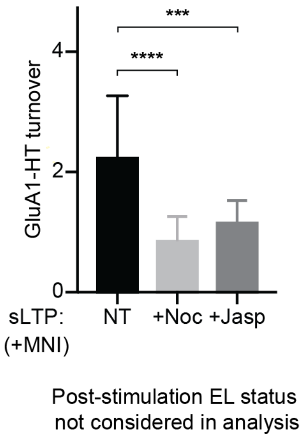
***
****
GluA1-HT turnover
4
2
0
sLTP:
(+MNI)
NT
+Noc
+Jasp
Post-stimulation EL status
not considered in analysis

**Figure S14. Nocodazole and jasplakinolide reduce sLTP-induced GluA1-HT turnover in spines lacking endosomes or lysosomes before stimulation, irrespective of status afterward.**

GluA1-HT turnover in response to structural long-term potentiation (sLTP) stimulation (+MNI) without chemical treatment (NT), in the presence of nocodazole (+Noc), or in the presence of jasplakinolide (+Jasp) near spines where mEmerald-tagged $Rab4^{+}$, $Rab7^{+}$, or $Rab11^{+}$ endosomes, or $LAMP1^{+}$ lysosomes are absent prior to stimulation, but EL status after stimulation is ignored. NT vs +Noc, ****p<0.0001; NT vs +Jasp, ***p=0.0003. Significance determined by Mann-Whitney test with Holm–Šídák correction across the comparisons shown in graph. N = 13–60 sLTP-stimulated spines for each condition pooled from 3–16 experiments each for Rab4, Rab7, Rab11, or LAMP1. For each experiment, images of the spine and EL marker were taken immediately before and after sLTP stimulation, while images of the EL marker and pulse-chase labeling were taken after the chase (see **Methods**). Turnover data for NT is a reanalysis of data from **Figure 3C**, +MNI $EL^{+}$ and $EL^{-}$ pooled together. Turnover data for +Noc is a reanalysis of **Figure 3E**, +Noc $EL^{-}$ plus the turnover data for one additional spine where an EL moved into position after sLTP stimulation (**Figure 3E**, top). Turnover data for +Jasp is a reanalysis of **Figure 3E**, +Jasp $EL^{-}$ plus the turnover data for 3 additional spines where ELs moved into position after sLTP stimulation (**Figure 3E**, top). This analysis shows nocodazole and jasplakinolide treatments reduce GluA1-HT turnover in spines lacking ELs prior to stimulation when the EL status after sLTP stimulation is not considered for analysis, motivating us to examine whether the EL status correlates with turnover when ELs are positioned before and after sLTP stimulation.

#### Figure S15

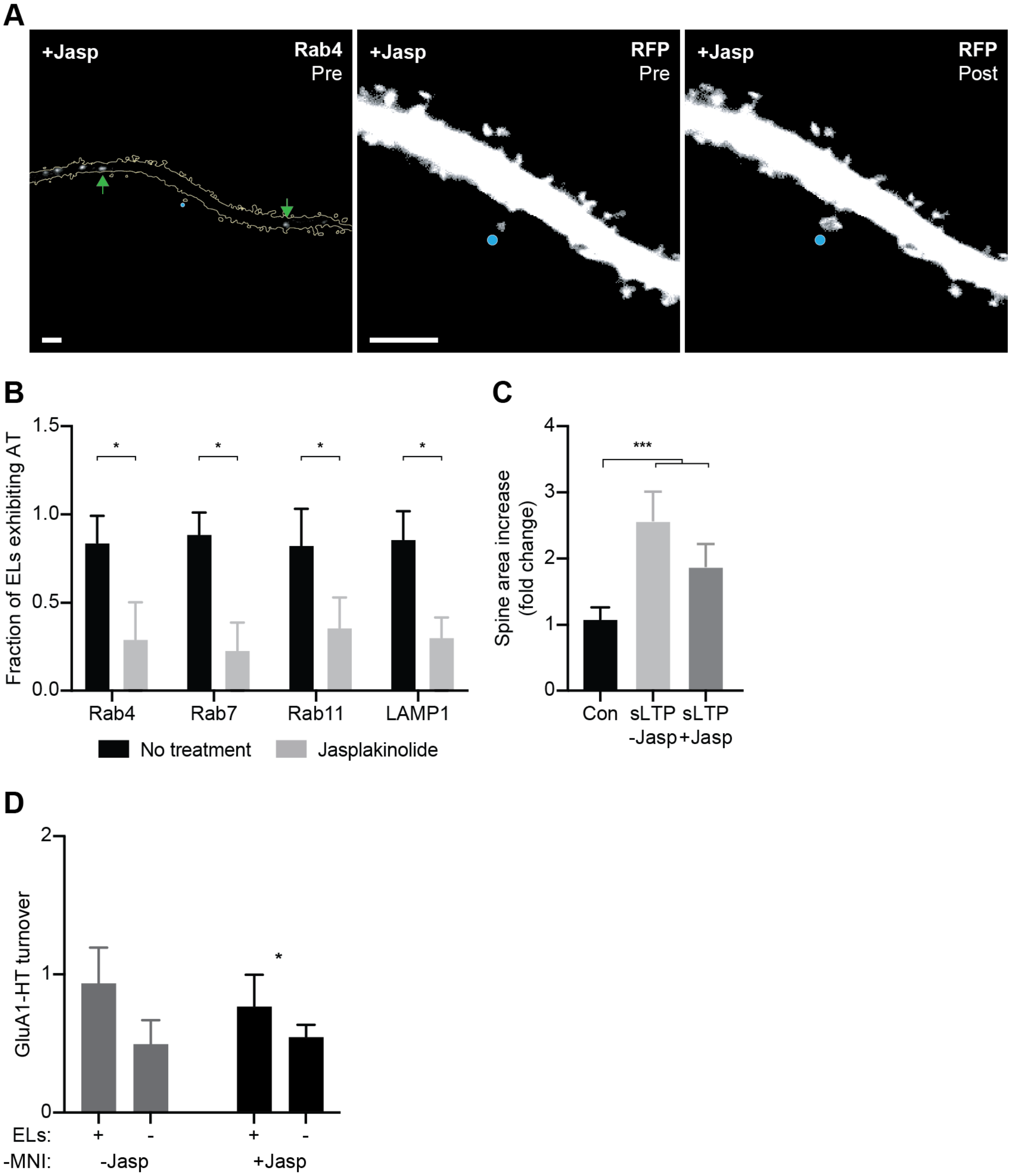

A
+Jasp
Rab4
Pre
+Jasp
RFP
Pre
+Jasp
RFP
Post
B
Fraction of ELs exhibiting AT
1.5
1.0
0.5
0.0
*
Rab4
Rab7
Rab11
LAMP1
No treatment
Jasplakinolide
C
Spine area increase (fold change)
4
3
2
1
0
***
Con
sLTP -Jasp
sLTP +Jasp
D
GluA1-HT turnover
2
1
0
*
ELs:
+
-
-MNI:
-Jasp
+Jasp

**Figure S15. Jasplakinolide treatment does not abolish spine enlargement in response to sLTP or inhibit GluA1-HT turnover in response to mock sLTP stimulation.**

(A) Rab4 (left): representative spinning disk confocal image of Rab4-mEmerald$^+$ (Rab4$^+$) endosomes in the presence of jasplakinolide (+Jasp) prior to structural long-term potentiation (sLTP) stimulation. Green arrows indicate Rab4$^+$ endosomes. RFP (right): dendritic spine depicted left (Rab4, left) expressing tdTomato pre-sLTP (Pre) and post-sLTP (Post) in the presence of jasplakinolide. Scale bars, 5 µm. Blue dot indicates MNI uncaging site.

(B) Fraction of mEmerald-tagged Rab4$^+$, Rab7$^+$, or Rab11$^+$ endosomes, or LAMP1$^+$ lysosomes (ELs) with active transport after treatment with jasplakinolide. Bars and error bars represent mean and standard deviation, respectively. Rab4, *p=0.0095; Rab7, *p=0.0095; Rab11, *p=0.0095; LAMP1, *p=0.0095. Significance determined by Mann-Whitney test with Holm–Šídák correction across four EL markers. N = 4–6 treatment experiments for each condition; for each experiment, the motion of ELs in a dendrite was imaged immediately after treatment. These data are a reproduction of the mock chemical long-term potentiation (cLTP) data and jasplakinolide treatment data from **Figure S4**.

(C) Increase in spine area (fold change) after sLTP in the presence of jasplakinolide. Bars and error bars represent mean and standard deviation, respectively. For control (Con), mock sLTP was performed without MNI (i.e., -MNI). Con vs sLTP -Jasp, ***p=0.0002; Con vs sLTP +Jasp, ***p=0.0006. Significance determined by Mann-Whitney test with Holm–Šídák correction across the comparisons shown in graph. N = 7–8 sLTP experiments for each condition; for each experiment, the target spine was imaged immediately before and after sLTP stimulation. The control and sLTP data are reproductions of the control and sLTP data from **Figure S13C**. A new neuronal culture was used for each sLTP experiment involving jasplakinolide treatment. We note that jasplakinolide reduces, but does not block, the fold change increase in spine area expansion in response to sLTP stimulation. This is likely because jasplakinolide itself stabilizes F-actin and leads to changes in spine morphology prior to sLTP.

(D) GluA1-HT turnover in response to mock sLTP stimulation (-MNI) in the absence or presence of jasplakinolide (-Jasp or +Jasp, respectively) on spines at which there are ELs present or absent (EL: +/-) near their bases before and after mock stimulation. Bars and error bars represent mean and standard deviation, respectively. -Jasp $EL^+$ vs +Jasp $EL^+$, p=0.1797; +Jasp $EL^+$ vs +Jasp $EL^-$, p=0.2491; +Jasp $EL^+$ vs +Jasp $EL^-$, *p=0.0245. Significance determined by Mann-Whitney test. N = 18–58 mock sLTP-stimulated spines for each condition pooled from 4–12 experiments each for Rab4, Rab7, Rab11, or LAMP1. For pulse-chase experiments analyzed in (D), we applied mock stimulation to dendritic spines that either did or did not have ELs near their bases prior to mock stimulation. EL positioning did not change in response to mock stimulation. For each experiment, images of the spine and EL marker were taken immediately before and after mock sLTP stimulation, while images of the EL marker and pulse-chase labeling were taken after the chase (see **Methods**). A new neuronal culture was used for each +Jasp experiment. -Jasp $EL^+$ turnover data is a reproduction of data from **Figure S13D**, -Noc $EL^+$. -Jasp $EL^-$ turnover data is a reproduction of turnover data from **Figure 3C**, -MNI $EL^-$.

#### Figure S16

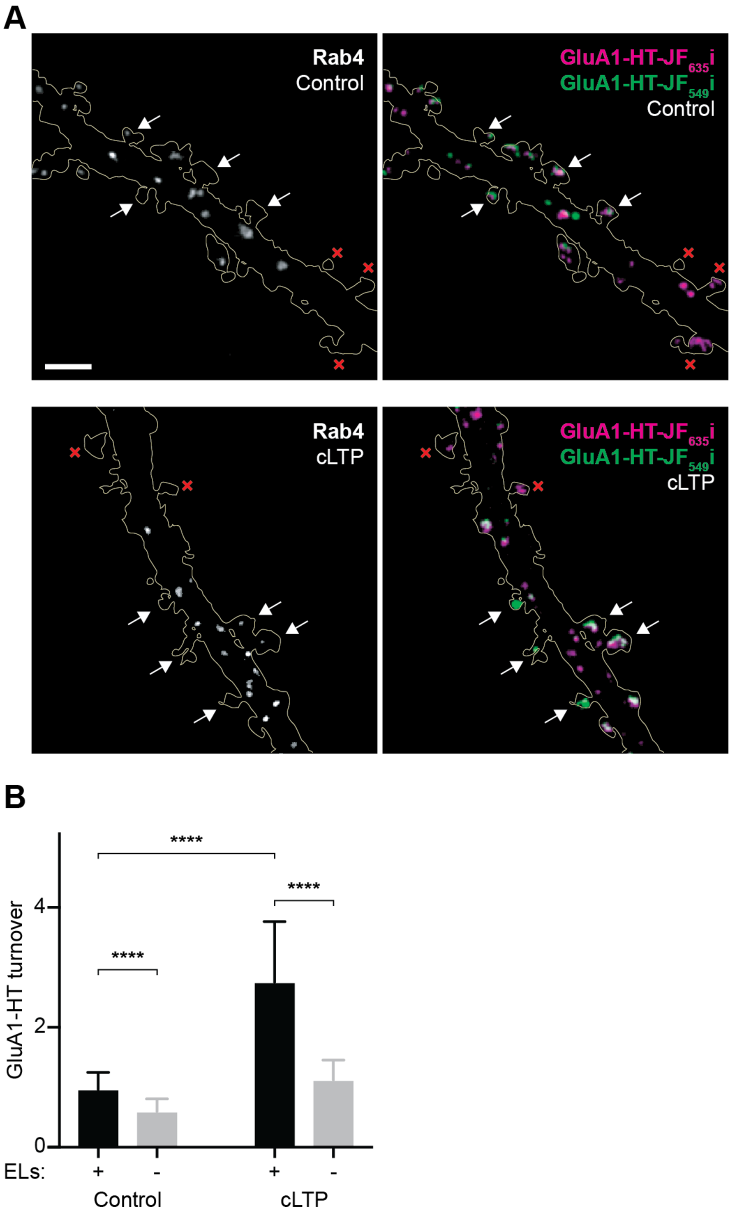
A
Rab4
Control
GluA1-HT-JF$_{635}$i
GluA1-HT-JF$_{549}$i
Control
Rab4
cLTP
GluA1-HT-JF$_{635}$i
GluA1-HT-JF$_{549}$i
cLTP
B
GluA1-HT turnover
0
2
4
****
****
****
ELs:
+
-
+
-
Control
cLTP

**Figure S16. Spines with endosome and lysosome positioning after cLTP have increased GluA1-HT turnover compared to spines that do not.**

(A) Rab4 (left): representative spinning disk confocal images of Rab4-mEmerald$^+$ (Rab4$^+$) endosomes after mock chemical long-term potentiation (cLTP) stimulation (i.e., stimulation buffer without glycine; Control, top) or cLTP stimulation (cLTP, bottom). Scale bar, 5 µm. GluA1-HT (right): images of GluA1-HT first labeled with $JF_{635}$i-HTL before treatment (pulse; magenta) and then labeled with $JF_{549}$i-HTL after treatment (chase; green). White arrows indicate the presence of a Rab4$^+$ endosome after stimulation. Red Xs indicate the absence of a Rab4$^+$ endosome after stimulation.

(B) GluA1-HT turnover in spines in which mEmerald-tagged Rab4$^+$, Rab7$^+$, or Rab11$^+$ endosomes, or LAMP1$^+$ lysosomes (ELs) are present (EL$^+$) or absent (EL$^-$) after either cLTP induction (cLTP) or mock cLTP induction (Control). Bars and error bars represent mean and standard deviation, respectively. Control, ****$p<0.0001$; cLTP, ****$p<0.0001$; Control EL$^+$ vs cLTP EL$^+$, ****$p<0.0001$. Significance determined by Mann-Whitney test with Holm–Šídák correction across the comparisons shown in graph. To confirm this result is not driven by non-independence among spines sharing the same neuronal culture, EL$^+$ vs EL$^-$ turnover under Control and cLTP conditions was additionally analyzed with a linear mixed-effects model including culture as a random intercept (Control, ****$p<0.0001$; cLTP, ****$p<0.0001$; see **Methods**). N = 26–41 spines for each condition (2–3 dendrites were imaged for each EL marker). At most two dendrites from two separate neurons were imaged from a single neuronal culture.

**Video 1. Timelapse images of Rab4$^+$ endosomes.**

Representative spinning disk confocal timelapse images of Rab4-mEmerald$^+$ (Rab4$^+$) endosomes in the dendrite of a cultured rat hippocampal neuron. Scale bar, 5 µm.

**Videos 2-5. Timelapse images of Rab4$^+$, Rab7$^+$, or Rab11$^+$ endosomes, or LAMP1$^+$ lysosomes in response to sLTP stimulation in neurons also expressing RFP.**

Representative spinning disk confocal or widefield timelapse images of Rab4-mEmerald$^+$ (Rab4$^+$; **Video 2**), Rab7-mEmerald$^+$ (Rab7$^+$; **Video 3**), or Rab11-mEmerald$^+$ (Rab11$^+$; **Video 4**) endosomes, or LAMP1-mEmerald$^+$ lysosomes (LAMP1$^+$; **Video 5**) in the dendrites of cultured rat hippocampal neurons before structural long-term potentiation (sLTP) stimulation (Pre, left) and after sLTP stimulation (Post, right). tdTomato (i.e., RFP) was used as a cytosolic marker to identify the dendritic shafts and spines, and to confirm spine volume expansion after sLTP stimulation. Scale bars, 5 µm. Blue dots indicate MNI uncaging site.

**Video 6. Timelapse images of Rab4$^+$ endosomes in response to mock sLTP stimulation in the absence of MNI.**

Representative spinning disk confocal timelapse images of Rab4-mEmerald$^+$ (Rab4$^+$) endosomes in the dendrite of a cultured rat hippocampal neuron before and after mock structural long-term potentiation (sLTP) stimulation in the absence of MNI (-MNI; Pre, left and Post, right, respectively). tdTomato (i.e., RFP) was used as a cytosolic marker to identify the dendritic shaft and spines. Scale bars, 5 µm. Blue dots indicate MNI uncaging site.

**Videos 7-10. Timelapse images of Rab4$^+$, Rab7$^+$, or Rab11$^+$ endosomes, or LAMP1$^+$ lysosomes in response to sLTP stimulation in neurons also expressing Tractin.**

Representative spinning disk confocal or widefield timelapse images of Rab4-mEmerald$^+$ (Rab4$^+$; **Video 7**), Rab7-mEmerald$^+$ (Rab7$^+$; **Video 8**), or Rab11-mEmerald$^+$ (Rab11$^+$; **Video 9**) endosomes, or LAMP1-mEmerald$^+$ lysosomes (LAMP1$^+$; **Video 10**) in the dendrites of cultured rat hippocampal neurons before sLTP stimulation (Pre, left) and after sLTP stimulation (Post, right). Tractin was expressed to define regions with increased actin polymerization in response to sLTP stimulation (see **Figure S8** for corresponding Tractin images). Scale bars, 5 µm. Blue dots indicate MNI uncaging site.
